# A conserved differentiation program facilitates inhibitory neuron production in the developing mouse and human cerebellum

**DOI:** 10.1101/2025.04.10.648162

**Authors:** Jens Bager Christensen, Alex P.A. Donovan, Marzieh Moradi, Giada Vanacore, Mohab Helmy, Adam J. Reid, Jimmy Tsz Hang Lee, Omer Ali Bayraktar, Andrea H. Brand, N. Sumru Bayin

## Abstract

Understanding the molecular mechanisms driving lineage decisions and differentiation during development is challenging in complex systems with a diverse progenitor pool, such as the mammalian cerebellum. Importantly, how different transcription factors cooperate to generate neural diversity and the gene regulatory mechanisms that drive neuron production, especially during the late stages of cerebellar development are poorly understood. Here, we used single cell RNA-sequencing (scRNA-seq) to investigate the developmental trajectories of Nestin-expressing progenitors (NEPs) in the neonatal mouse cerebellum. We identified FOXO1 as a key regulator of NEP-to-inhibitory neuron differentiation, acting directly downstream of ASCL1. Genome occupancy and functional experiments using primary NEPs showed that both ASCL1 and FOXO1 regulate neurogenesis genes during differentiation while independently regulating proliferation and survival, respectively. Furthermore, we demonstrated that WNT signalling promotes the transition from an ASCL1^+^ to a FOXO1^+^ cellular state. Finally, we showed that the role of WNT signalling in promoting neuron production via FOXO1 is conserved in primary human NEPs. By resolving how cerebellar interneurons differentiate, our findings could have implications for cerebellar disorders such as spinocerebellar ataxia, where cerebellar interneurons are overproduced.

## Introduction

How neural stem cells generate diverse cell types with intricate cytoarchitecture in the brain remains a fundamental yet unresolved question in developmental biology. While fate mapping and clonal analyses are instrumental in resolving how distinct lineages are generated during nervous system development, the molecular mechanisms that drive lineage decisions and differentiation remain poorly understood, especially outside the forebrain. Single-cell genomic technologies have enabled the capture of transient cell states during lineage commitment and differentiation, providing the opportunity to map the dynamics of the gene regulatory networks (GRNs) driving these processes.

The cerebellum is a key hindbrain structure, crucial for motor coordination, as well as cognitive and social behaviours (Caligiore et al., 2017; Fatemi et al., 2012; Koziol et al., 2014; Lackey et al., 2018). Having co-evolved with the neocortex (Herculano-Houzel, 2010), the cerebellum has significantly expanded in mammals, housing ∼60% and ∼80% of the neurons in the mouse and human brains, respectively (Azevedo et al., 2009; Gill and Sillitoe, 2019; Herculano-Houzel et al., 2006). Although the cerebellar primordium is established at embryonic day (E) 9 in mice (Leto et al., 2016), the majority of its growth occurs postnatally. The postnatal cerebellar progenitors continue to proliferate and differentiate up to two weeks after birth in mice and six months in humans (Leto et al., 2016; Rakic and Sidman, 1970). Due to this protracted development, the cerebellum is particularly vulnerable to stress and damage around birth. Indeed, cerebellar injury at birth is the second leading risk factor for autism spectrum disorders (Wang et al., 2014). Furthermore, increasing evidence suggests that spinocerebellar ataxias manifest as developmental abnormalities in various cerebellar cell types, including altered progenitor cell behaviour and postnatal inhibitory interneuron overproduction (Becker et al., 2009; Edamakanti et al., 2018; Luttik et al., 2022). Therefore, dissecting the molecular mechanisms that regulate postnatal cerebellum development is critical to understanding disease pathophysiology and identifying potential therapeutic interventions.

The postnatal development of the cerebellum is orchestrated by the coordinated action of lineage-restricted progenitors derived from two molecularly distinct embryonic germinal zones: the rhombic lip and the ventricular zone (VZ) (Joyner and Bayin, 2022; Leto et al., 2016; Wizeman et al., 2019). Rhombic lip-derived granule cell progenitors (GCPs) proliferate in the external granule layer and differentiate into the excitatory granule cells that migrate inwards to the internal granule layer in an outside-in manner (Machold and Fishell, 2005; Wang et al., 2005; Wingate and Hatten, 1999). On the other hand, VZ-derived *nestin*-expressing progenitors (NEPs) consist of multiple subtypes and give rise to molecular layer inhibitory neurons, Bergmann glia and astrocytes (Bayin et al., 2021; Lee et al., 2005; Milosevic and Goldman, 2004; Parmigiani et al., 2015). The *Hopx*-expressing gliogenic NEPs (*Hopx*-NEPs) are further divided into two subtypes based on their locations: those in the lobule white matter (WM) generate astrocytes, while those in the Bergmann glia layer (BgL) give rise to both astrocytes and Bergmann glia (Bayin et al., 2021; Cerrato et al., 2018). The neurogenic NEPs that express the basic helix loop helix proneural transcription factor *Ascl1* (*Ascl1*-NEPs) reside in the lobule WM of the postnatal cerebellum. The *Ascl1*-NEPs proliferate in the WM and generate the molecular layer inhibitory interneurons, basket and stellate cells (Leto et al., 2009). Once born in the lobule WM, the inhibitory neurons migrate radially to the molecular layer in an inside-out manner (Leto et al., 2006; Leto et al., 2009; Leto et al., 2016; Sudarov et al., 2011; Zhang and Goldman, 1996). Furthermore, lineage tracing studies have demonstrated that some NEPs in the WM are bipotent, generating both astrocytes and interneurons during early postnatal development (Bayin et al., 2021; Cerrato et al., 2018). Finally, a small subset of GCPs also express the NEP markers *Sox2* and *Nes* (Selvadurai et al., 2020; Vanner et al., 2014), however, whether these are derived from the rhombic lip or VZ remains unclear. While the molecular mechanisms that regulate GCP proliferation and differentiation have been extensively studied due to their role in some medulloblastomas (Cheng et al., 2020; Dahmane and Altaba, 1999; Gold et al., 2024; Miyata et al., 1999), the molecular mechanisms that govern NEP subpopulations remain largely unknown.

Cellular identities and the differentiation programmes required to achieve them are established by the combinatorial and context-dependent action of pleiotropic transcription factors. A recent analysis revealed that multiple transcription factors cooperate to mediate GABAergic differentiation in cerebral interneurons, highlighting the importance of transcription factor networks in development (Catta-Preta et al., 2024). However, the gene regulatory networks that drive molecular layer inhibitory neuron production in the cerebellum are not well understood, although ASCL1 is shown to be required for their differentiation (Grimaldi et al., 2009; Sudarov et al., 2011). ASCL1 is a pioneer proneural transcription factor that drives neurogenesis (Castro et al., 2006; Raposo et al., 2015; Woods et al., 2022) during development (Casarosa et al., 1999) and in direct reprogramming (Vierbuchen et al., 2010). A candidate family of proteins that could cooperate with ASCL1 is the forkhead-box transcription factor family subgroup O (FOXO) proteins. They play a crucial role in stem cell homoeostasis (Paik et al., 2009), cellular metabolism(Bastie et al., 2005; Puigserver et al., 2003) and apoptosis (Brunet et al., 2004; Papadia et al., 2008; Yuan et al., 2008). While some studies show that FOXO genes are important for neural stem cell homeostasis and promote stemness by inhibiting differentiation (Kim et al., 2015; Paik et al., 2009; Webb et al., 2013), others have suggested that they regulate autophagy, migration and maturation (De La Torre-Ubieta et al., 2010; Huynh et al., 2011; Schäffner et al., 2018) in developing neurons as well. Finally, various ChIP-seq data suggest that ASCL1 and FOXO motifs co-occur in neural and brain tumour stem cells, suggesting overlapping roles (Park et al., 2017; Webb et al., 2013). While previous studies have shown that FOXO3 is a negative regulator of ASCL1-dependent neurogenesis (Webb et al., 2013), the functional relationship between FOXO proteins and ASCL1 remains largely unknown.

In this study, we used scRNA-seq, primary mouse and human NEP cultures and genetic induced fate mapping to investigate the molecular drivers of NEP-to-inhibitory neuron differentiation in the developing postnatal cerebellum. We demonstrated that during differentiation, ASCL1 and WNT signalling upregulate FOXO1, which in turn promotes neural differentiation. Analysis of genome occupancy showed that ASCL1 and FOXO1 regulate a common set of neurogenic genes but also exhibit temporally divergent functions; ASCL1 promotes the proliferation of *Ascl1*-NEPs, whereas FOXO1 supports cell survival during differentiation. Finally, we demonstrated that the molecular mechanisms governing NEP-to-inhibitory neuron differentiation in mouse NEPs are also conserved in humans. Collectively, this study establishes core components of the GRN that drive NEP-to-inhibitory neuron differentiation in the developing mouse and human cerebellum.

## Results

### Iterative subclustering reveals cellular states during NEP-to-inhibitory neuron differentiation during postnatal development

To identify key cellular states and the molecular mechanisms that drive NEP-to-inhibitory neuron differentiation during postnatal cerebellum development, we analysed previously generated scRNA-seq data of FACS-isolated CFP^+^ cells from postnatal day (P) 1, 2, 3 and 5 *Nes-Cfp/+* mouse cerebella (Pakula et al., 2025) (Fig. 1A-D). Following quality control, 4872 cells were clustered into 11 clusters to identify distinct NEP subtypes and cellular states during the first 5 days after birth (Fig. 1E and S1A). Clusters were annotated using established lineage markers and highlighted the presence of the expected NEP subtypes, their immediate progenies likely captured as a result of CFP perdurance, and other *Nes^+^*populations as previously observed (Pakula et al., 2025). We identified *Hopx-*NEPs (clusters 0 and 6), *Ascl1*-NEPs (cluster 1), astrocytes (cluster 4, *Slc6a11*^+^) and immature inhibitory neurons (cluster 3, *Pax2*^+^). In addition, CFP^+^ granule cell progenitors (clusters 2, 5 and 8, *Atoh1*^+^/*Barlh1*^+^), oligodendrocyte progenitors and oligodendrocytes (cluster 10, *Olig2*^+^/*Pdgfra*^+^), ependymal cells (cluster 7, *Foxj1*^+^), mesenchymal cells (cluster 9, *Vtn*^+^) and microglia (cluster 11, *Cx3cr1*^+^) were detected in various proportions across different postnatal days (Figs 1E, S1A-C, Table S1).

**Fig. 1.**
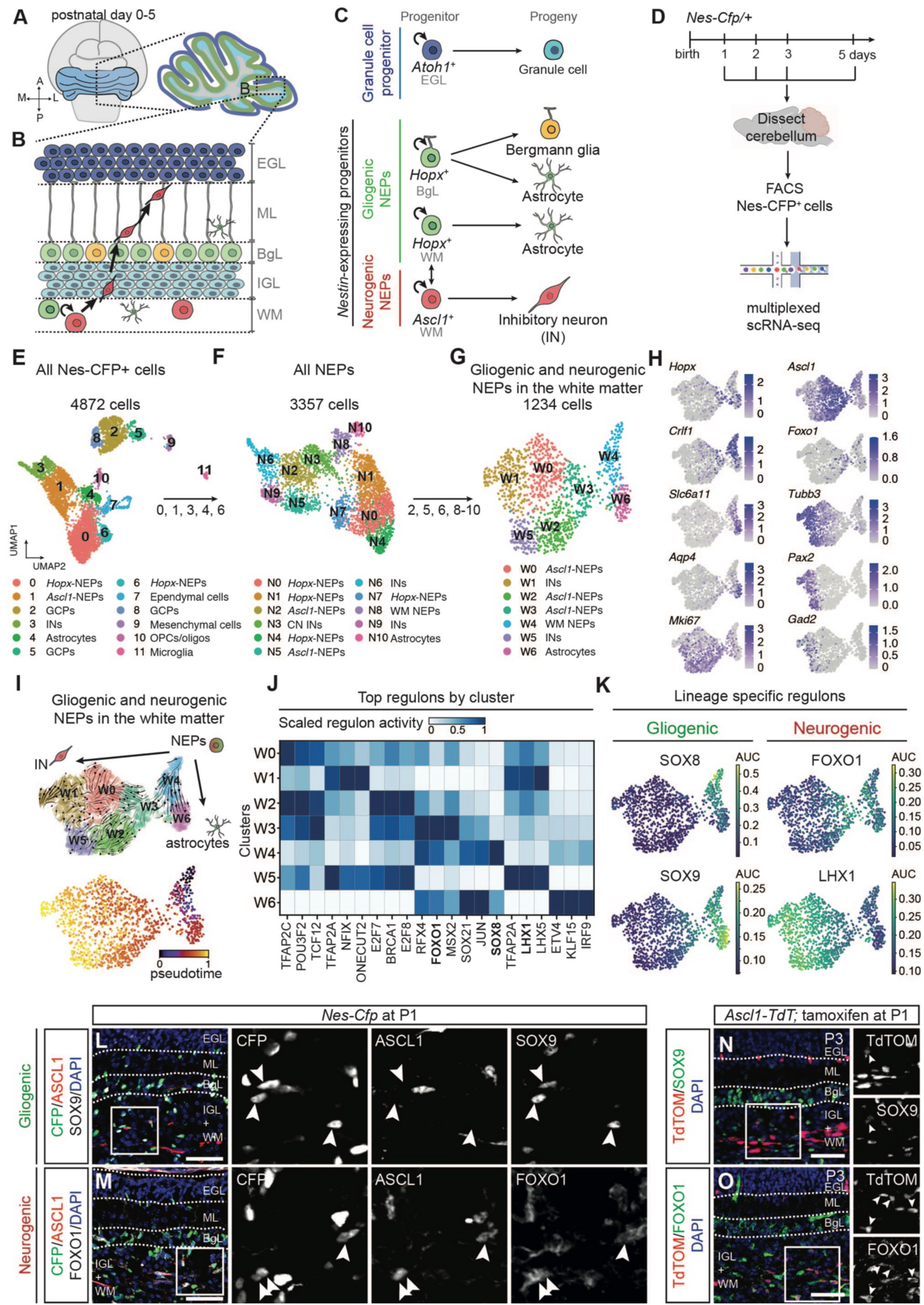
scRNA-seq captures the molecular diversity of NEPs and reveals the regulatory mechanisms that drive the lineage decisions and differentiation during postnatal mouse cerebellum development. (A-C) Schematics showing the postnatal cerebellar progenitors and their lineages. (D) Experimental design. (E-G) Uniform Manifold Approximation and Projection (UMAP) visualisation of all Nes-CFP^+^ cells (4872 cells, E), all NEPs (3357 cells, F) and the gliogenic and neurogenic NEPs in the WM (1234 cells, G) labelled by cluster identity (Table S1 and Fig. S1). (H) UMAP projection of normalised expression levels of known marker genes (gliogenic-NEPs: *Hopx*; astrocytes: *Slc6a11*, *Aqpr4*; neurogenic-NEPs: *Ascl1*; immature inhibitory neurons: *Tubb3, Pax2, Gad2*), genes that were significantly enriched in W3 (*Foxo1)* and W4 (*Crlfl1*), and proliferation markers (*mKi67*). (I) UMAP projection of cellular trajectories (top) or pseudotime (bottom) within the WM NEPs, computed using scVelo. (J) Heatmap of the normalised activity scores of the top 3 cluster-specific regulons identified by pySCENIC (Table S2). (K) UMAP projection of regulon activity on WM NEPs. (L-M) Immunofluorescent analysis of P1 *Nes*-*Cfp*/+ cerebella for CFP, ASCL1 and SOX9 (L) or FOXO1 (M). Arrowheads in (L) show CFP^+^/ASCL1^-^/SOX9^+^ cells, and in (M) show CFP^+^/ASCL1^+^/FOXO1^+^ triple positive cells. (N-O) Immunofluorescent analysis of P3 *Ascl1-TdT* pups that were given Tamoxifen at P1. Arrowheads indicate TdTOM^+^/SOX9^+^ (N) or TdTOM^+^/FOXO1^+^ (O) cells. Scale bars: 50 µm. OPC/oligo: oligodendrocyte progenitor cells/oligodendrocytes, INs: inhibitory neurons, CN: cerebellar nuclei, EGL: external granule layer, IGL: internal granule layer, ML: molecular layer, BgL: Bergmann glia layer, WM: white matter.

To increase the resolution of distinct NEP cellular states and to identify mechanisms of NEP-to-inhibitory neuron differentiation, we performed iterative subclustering. First, we subclustered all NEPs from the initial dataset (clusters 0, 1, 3, 4 and 6, Fig. 1E). Clustering of 3357 NEPs showed increased molecular diversity highlighted by the presence of ten clusters (Figs 1F and S1D-F, Table S1). BgL *Hopx*-NEPs (clusters N0, N1, N4 and N7, *Gdf10*^+^), WM *Hopx*- and *Ascl1*-NEPs (i.e. *Hopx*-(cluster N8) and *Ascl1*-NEPs (clusters N2 and N5)) were identified along with their immediate progenies, *Slc6a11*^+^ astrocytes (cluster N10) and immature *Pax2*^+^ inhibitory neurons (clusters N6 and N9). Finally, cluster N3 showed enrichment for *Pvalb* expression (Fig. S1F). Based on Allen brain atlas RNA *in situ* hybridisation data at P4, *Pvalb* is expressed in the cerebellar nuclei in the deep white matter (Fig. S1G), and thus, cluster N3 was excluded from further analyses.

Next, we aimed to identify WM NEPs in the cerebellar cortex, which generate the molecular layer inhibitory neurons and astrocytes. To this end, we further subclustered specifically the 1234 gliogenic and neurogenic NEPs in the WM (clusters N2, N5, N6, N8-10) (Figs 1F-G, S1H-I and Table S1). This analysis revealed seven distinct clusters, highlighting WM NEP subtypes, one of which was a potential bipotent NEP state (cluster W4, *Ascl1*^+^/*Hopx*^low^/*Crlf1*^+^) consistent with the previous fate mapping and clonal analyses(Bayin et al., 2021; Cerrato et al., 2018) (Fig. 1G-H). While only one cluster of astrocytes was detected (W6, *Slc6a*11^+^), likely due to their later production compared to neurons, we observed 5 clusters (W0-3 and W5, *Ascl1*^+^ and/or *Tubb3^+^*) with neural identity, differentiated by expression of transcription factors such as *Foxo1* (W3), and/or their proliferation (W2, W3, W5, *mKi67^+^*) and maturation (immature inhibitory neurons, W0-1, W5, *Pax2^+^, Gad1/2^+^*) state (Fig. 1G-H, S1H-I and Table S1). In summary, iterative subclustering of scRNA-seq data resolved distinct cellular states within known NEP subtypes during early postnatal cerebellum development.

### Distinct gene regulatory networks drive NEP lineage progression

To identify the transcriptional mechanisms that drive the differentiation of WM NEPs into inhibitory neurons or astrocytes, we first performed RNA velocity analysis using scVelo (Bergen et al., 2020). As expected, bipotent cells were positioned at the top of the WM NEP lineage hierarchy, leading to two trajectories, either towards inhibitory neurons (*Gad2*^+^) via a *Foxo1*+ state, or astrocytes (*Slc6a11*^+^) (Fig. 1I). Interestingly, various intermediate states were detected along the inhibitory neuron differentiation path, including a subset with persistent *Ascl1* and *mKi67* expression (*Ascl1*-NEPs) (Figs 1H-I and S1J). To further explore the molecular drivers of glial and neuronal differentiation, we performed *in silico* GRN reconstruction using pySCENIC(Van de Sande et al., 2020). This analysis computes an activity score for regulons based on the coordinated expression of a transcription factor and its known target genes. Analysis of the top enriched regulons for each cluster revealed both lineage and stage-specific transcription factors that could be involved in maintaining stemness, bipotency, lineage decisions or differentiation (Fig. 1J-K and Table S2). For example, the SOX8 regulon was specifically active in bipotent NEPs (cluster W4), whereas LHX1/5 and SOX9 regulons were highly active in immature inhibitory neurons (clusters W1 and W5) and astrocytes (cluster W6), respectively (Figs 1J-K and Table S2). These findings are in line with previous reports from other brain regions (Kang et al., 2012; Pillai et al., 2007; Seto et al., 2014; Stolt et al., 2003; Sun et al., 2017; Vong et al., 2015). Finally, we hypothesised that the regulons enriched in cluster W3 could play crucial roles during neural differentiation. Among these regulons, FOXO1 was identified as one of the specific regulons in cluster W3 (Fig. 1J, K), overlapping with its high expression in that cluster (Fig. 1H).

To ascertain potential functional roles for these transcription factors, we then performed *in silico* gene overexpression and knockout using CellOracle (Kamimoto et al., 2023) (Fig. S1K-P). Knockout of *Sox8* did not significantly alter the computed cellular trajectories, whereas overexpression of *Sox8* was predicted to promote stemness over differentiation in the bipotent NEPs (Fig. S1L). On the other hand, *in silico* knockout of *Sox9* promoted, whereas knockout of *Ascl1* and *Lhx1* inhibited, neuron differentiation (Fig. S1M-O). In contrast, the overexpression of these transcription factors had opposite effects (Fig. S1M-O), in line with their known roles during neural and glial differentiation (Pillai et al., 2007; Stolt et al., 2003; Sudarov et al., 2011; Vong et al., 2015). *In silico* knockout and overexpression of *Foxo1* inhibited and promoted neuronal differentiation, respectively (Fig. S1P), supporting its potential role during postnatal cerebellar neurogenesis.

To validate the expression of transcription factors identified in our scRNA-seq analyses, we performed immunofluorescent staining on sections obtained from P1 *Nes-Cfp/+* cerebella. Immunofluorescent analysis of SOX9, which was enriched in NEPs (W4) and astrocytes (W6), showed expression in a subset of CFP^+^ cells in the WM, albeit mutually exclusive from ASCL1 (Fig. 1L). In contrast, we observed nuclear FOXO1 in the CFP^+^ cells in the WM, some of which were also ASCL1^+^, and in endothelial cells, as previously reported(Hosaka et al., 2004) (Fig. 1M). Similarly, fate mapping using *Ascl1^CreERT2/+^;R26^lox-STOP-lox-TdTom/+^*(*Ascl1-TdT*) animals, where tamoxifen was administered at P1 and analysed 2 days after, revealed only rare SOX9^+^/TdTOM^+^ cells, while 19.94% ± 0.11% (n=3) of TdTOM^+^ cells in the WM had nuclear FOXO1 expression (Fig. 1N-O). FOXO1^+^/TdTOM^+^ cells were restricted to the WM and were not observed in the TdTOM^+^ cells that had migrated to outer layers, highlighting the transient nature of *Foxo1* expression during differentiation. Interestingly, although *Foxo1* mRNA was present in the gliogenic-NEPs in the BgL, immunofluorescent staining on sections from P1 and P5 *Nes-Cfp/+* cerebella showed cytoplasmic localisation, suggesting limited transcriptional activity in this population (Fig. S1Q-T). In summary, using scRNA-seq, we identified GRNs that regulate WM NEP lineage decisions towards astrocytes or neurons, and underscore *Foxo1* as a potential regulator of postnatal cerebellar inhibitory neuron production.

### ASCL1 precedes FOXO1 during postnatal NEP-to-inhibitory neuron differentiation *in vitro*

The scRNA-seq and histological analyses suggested that ASCL1 precedes FOXO1 expression during NEP-to-inhibitory neuron differentiation (Fig. 1H-I, 1K, 1N-O). To further resolve the temporal dynamics of ASCL1 and FOXO1 expression during postnatal NEP-to-inhibitory neuron differentiation and to investigate the molecular drivers of differentiation, we adopted an *in vitro* approach. Primary NEPs were isolated from P1 cerebella and cultured as neurospheres in suspension with 20ng/mL EGF and FGF2, similar to other neural stem cell cultures (Capela and Temple, 2002; Rietze et al., 2001). NEPs could then be dissociated and differentiated as adherent cultures into neurons and glia by withdrawing growth factors and treating with 2% fetal bovine serum (FBS) (Fig. 2A). Almost all cells in the neurospheres were SOX2^+^ and expressed either ASCL1 or HOPX, demonstrating the presence of both *Ascl1*- and *Hopx*-NEPs (Fig. 2B). Upon differentiation, the number of TUJ1^+^ neurons gradually increased over time and plateaued starting at day 10 (Fig. 2C-F). At the end of the 14-day differentiation period, we observed mixed cultures with GFAP^+^ and TUJ1^+^ cells (14.74 ± 2.81%, n=9) (Fig. 2F). We observed that some of the TUJ1^+^ cells were also GAD1/2^+^, confirming their inhibitory neuron identity (Fig. 2E).

**Fig. 2.**
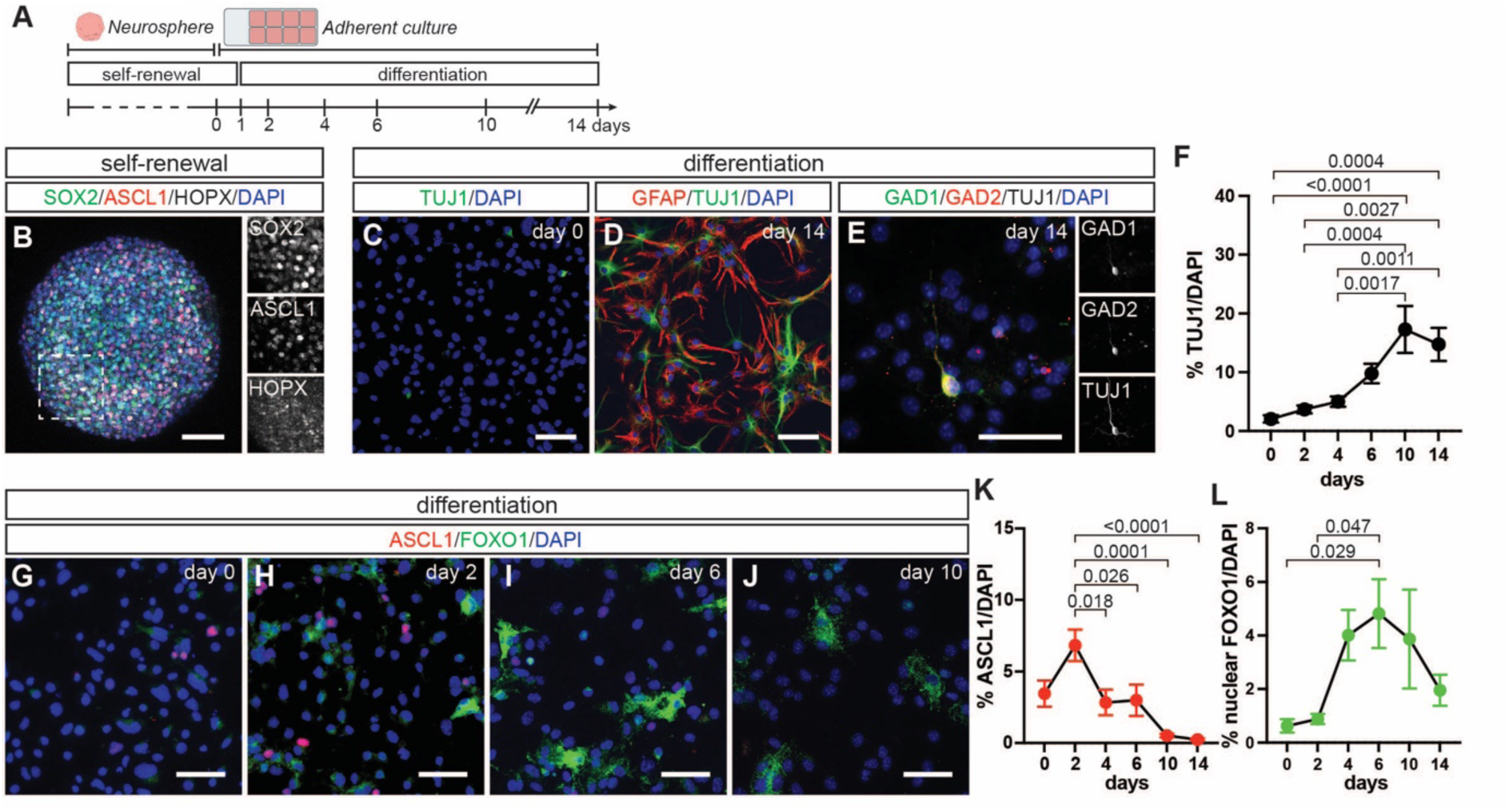
NEP-to-inhibitory neuron differentiation *in vitro* reveals the temporal dynamics of ASCL1 and FOXO1. (A) Experimental design. (B) Immunofluorescent analysis of primary NEP neurospheres for SOX2 (pan-NEP), ASCL1 (neurogenic-NEP) and HOPX (gliogenic-NEP). (C-E) Immunofluorescent analysis of adherent NEP cultures at day 0 (C) and 14 (D-E) of differentiation. (F) Quantification of TUJ1^+^ cells during *in vitro* differentiation (one-way ANOVA, F_(5,46)_=9.439, p<0.0001, n=9 (except 10, n=7)). (G-J) Immunofluorescent analysis for ASCL1 and FOXO1 during differentiation. Representative images from days 0 (G), 2 (H), 6 (I) and 10 (J) are shown. (K-L) Quantification of ASCL1^+^ (K) and nuclear FOXO1^+^ (L) cells during differentiation (one-way ANOVA, L: F_(5,46)_=7.720, p<0.0001, n=9 (except 10, n=7). K: F_(5,46)_=3.575, p=0.0082, n=9 (except 10, n =7)). Graphs show mean ± s.e.m. and significant p-values for multiple comparisons are shown in the figure. Scale bars: 50 µm, except for E (25 µm).

Next, we analysed the temporal dynamics of ASCL1 and FOXO1 expression during *in vitro* differentiation (Fig. 2G-L). At day 0 of differentiation, a small proportion of cells were ASCL1^+^ (3.46% ± 0.91%, n=9), likely representing the *Ascl1-expressing* neurogenic NEPs, whereas nuclear FOXO1 cells were minimal (0.62% ± 0.25%, n=9) (Fig. 2G, K-L). During differentiation, we first observed a 1.98 ± 0.32-fold (n=9) increase in the proportion of ASCL1^+^ cells on day 2 compared to day 0, which gradually decreased to 0.25% ± 0.12% (n=9) at 14 days (Fig. 2K). On the other hand, the proportion of cells with nuclear FOXO1 increased after the peak of ASCL1^+^ cells (day 2), starting at around day 4 of differentiation (7.81 ± 1.84-fold compared to day 0, n=9) and maintaining similar levels until day 10 with a downwards trend at the end of differentiation at day 14 (Fig. 2L). Collectively, these results demonstrate that ASCL1 precedes FOXO1 during *in vitro* differentiation of NEPs into inhibitory neurons, in agreement with the scRNA-seq and *Ascl1-Tdt* fate mapping results.

### ASCL1 and FOXO1 both drive neurogenesis *in vitro*

*In vivo* studies have shown that ASCL1 is required for inhibitory neuron production in the developing cerebellum (Sudarov et al., 2011). However, the role of FOXO1 during NEP-to-inhibitory neuron differentiation and whether ASCL1 and FOXO1 interact during differentiation remain to be understood. To further resolve their functions, we next performed gain- and loss-of-function studies in our *in vitro* culture paradigm (Fig. 3A). To test whether ASCL1 and FOXO1 alone can promote neurogenesis, we used a doxycycline (DOX)-inducible lentiviral system to overexpress them in a temporally restricted manner (Fig. 3A). We established a stable rTTA-NEP cell line, which was then infected with lentiviruses that overexpressed either ASCL1 (ASCL1 OE), FOXO1 (FOXO1 OE) or green fluorescent protein (EGFP OE) under the control of a TET-responsive promoter (Liu et al., 2018; Panciera et al., 2016) (Fig. 3A). Quantification of EGFP^+^ cells in the EGFP OE NEPs demonstrated transduction efficiencies of 30.33% ± 2.38% (n=5) at day 4 and 33.88% ± 1.61% (n=5) at day 10 of differentiation (Figs 3B-C and S2E). Unfortunately, the TET-inducible promoter was leaky in primary NEPs, and we detected similar frequencies of EGFP^+^ cells with or without DOX treatment (Fig. S2A and S2E). Therefore, we focused on DOX-treated cells for downstream analyses. No significant changes were observed in the parameters assessed between control cells and cells treated with DOX or vehicle (Fig. S2A-H). However, the level of overexpression was higher with DOX (Fig. S2I-K).

**Fig. 3.**
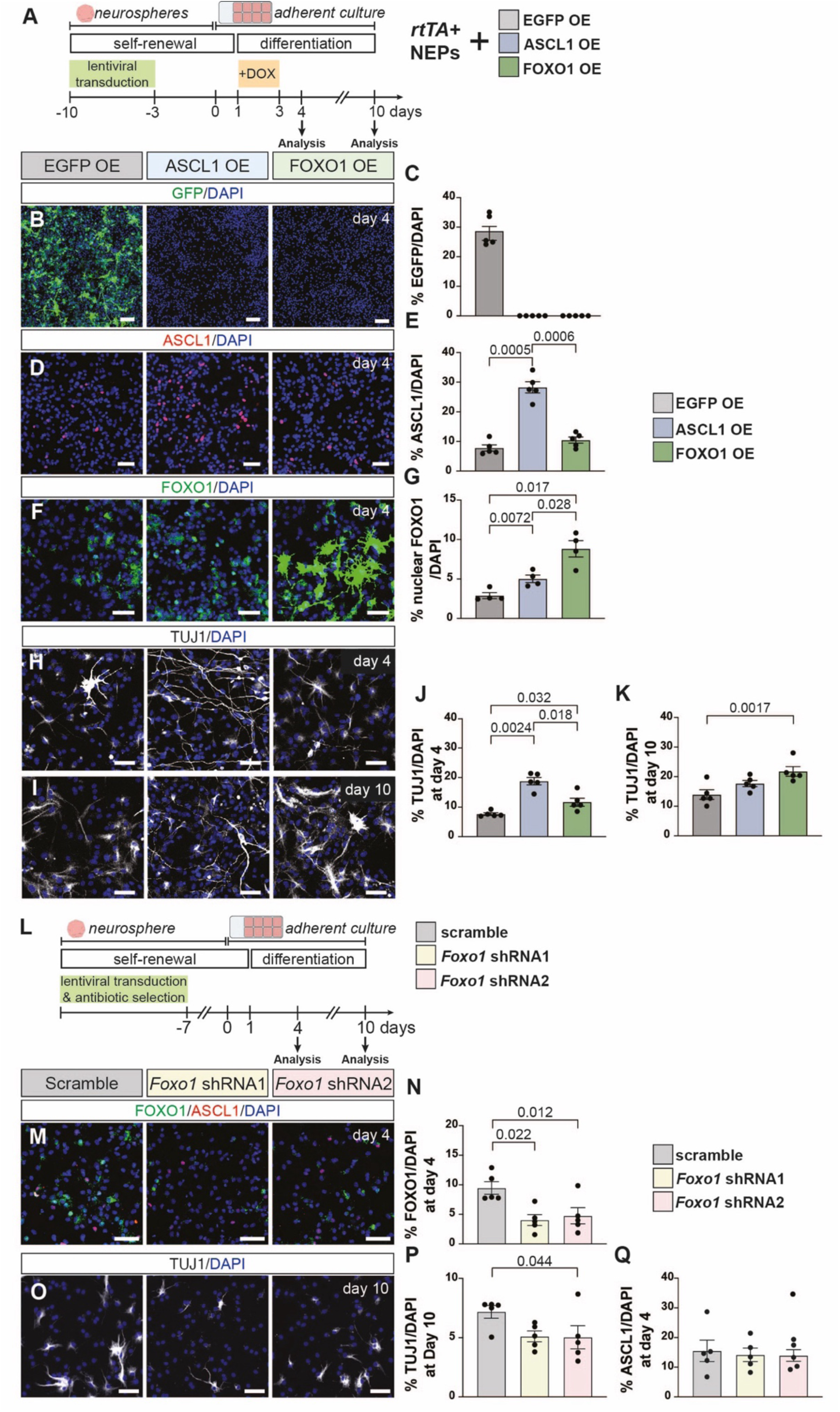
FOXO1 is downstream of ASCL1 and promotes neuron production *in vitro*. (A) Experimental design. (B-G) Immunofluorescent analysis of differentiating cells at day 4 that are overexpressing EGFP (EGFP OE), ASCL1 (ASCL1 OE) or FOXO1 (FOXO1 OE). EGFP^+^ (B-C), ASCL1^+^ (D-E) or FOXO1^+^ (F-G) cells were quantified (one-way ANOVA, A: F_(1,4)_=149, p=0.0003, n=5; B: F_(1,5)_=138.9, p<0.0001, n=5; C: F(_1,3_)=35.83, p=0.0087, n=4). (H-I) Immunofluorescent analysis of differentiating cells at day 4 (H) or 10 (I) EGFP OE, ASCL1 OE and FOXO1 OE cultures. (J-K) TUJ1^+^ cells were quantified on day 4 (J) and 10 (K) (one-way ANOVA, J: F_(1,6)_=39.25, p=0.0002, n=5; K: F_(1,5)_=17.37, p=0.0044, n=5). (L) Experimental design. (M) Immunofluorescent analysis of cultures overexpressing scramble, *Foxo1* shRNA1 or *Foxo1* shRNA2 for FOXO1 and ASCL1 at day 4 of differentiation. (N) Quantification of FOXO1^+^ cells at day 4 in the scramble, *Foxo1* shRNA1 or *Foxo1* shRNA2 expressing cells (n=5, paired Student’s t-test, compared to scrambled). (O) Immunofluorescent analysis of cells that are overexpressing scramble, *Foxo1* shRNA1 or *Foxo1* shRNA2 for TUJ1 at day 10. (P) Quantification of TUJ1^+^ cells at day 10 (P) or ASCL1^+^ cells at day 4 (Q) (n=5, paired Student’s t-test, compared to scrambled). Representative images are shown. Graphs show mean ± s.e.m. and significant p-values for multiple comparisons are shown in the figure. Scale bars: 50 µm.

First, we confirmed overexpression by quantifying the percentage of ASCL1^+^ and nuclear FOXO1^+^ cells in each overexpression condition during NEP differentiation. On day 4, all conditions showed significant upregulation of the respective protein (Figs 3B-G). Interestingly, ASCL1 OE also showed a significant increase in nuclear FOXO1^+^ cells (Fig. 3G), suggesting that ASCL1 promotes FOXO1 expression. Next, we addressed the effect of overexpression on neurogenesis by quantifying the number of neurons (TUJ1^+^) in each condition during differentiation (Fig. 3H-K). We observed a significant increase in the proportion of neurons upon both ASCL1 OE and FOXO1 OE at day 4, with the fold-change relative to EGFP OE being higher in ASCL1 OE compared to FOXO1 OE (2.46 ± 0.16-fold vs. 1.54 ± 0.18-fold, n=5, Fig. 3J). Surprisingly, while the FOXO1 OE condition continued to have significantly more TUJ1^+^ cells at day 10 of differentiation, the ASCL1 OE did not (Fig. 3K). Furthermore, the TUJ1^+^ cells in the ASCL1 OE condition had longer processes compared to the EGFP OE and FOXO1 OE conditions both at day 4 and 10, suggesting that sustained overexpression of these factors that function at the early stages of differentiation may affect neural maturation or even lead to cell death.

To test whether FOXO1 is required for NEP-to-inhibitory neuron differentiation, we performed lentiviral knockdown using shRNAs against *Foxo1* during *in vitro* differentiation. Stable primary NEP lines expressing either shRNAs against *Foxo1* or a scrambled shRNA as control were generated and then differentiated (Fig. 3L). Quantification of FOXO1^+^ cell numbers showed a significant reduction in the percentage of FOXO1^+^ cells (FOXO1 shRNA1: 0.43 ± 0.10-fold, FOXO1 shRNA2: 0.50 ± 0.14-fold, n=5), confirming the knockdown (Fig. 4M-N). FOXO1 knockdown led to a significant reduction in the percentage of TUJ1^+^ cells at day 10 of differentiation (shRNA1: 0.71 ± 0.06-fold and shRNA2: 0.70 ± 0.14-fold, n=5), while the number of ASCL1^+^ cells did not change at day 4 (Fig. 3O-Q). Together, this data demonstrates that both ASCL1 and FOXO1 promote the production of neurons *in vitro* but to different extents and suggests that ASCL1 regulates FOXO1 expression during the differentiation of NEPs to inhibitory neurons.

**Fig. 4.**
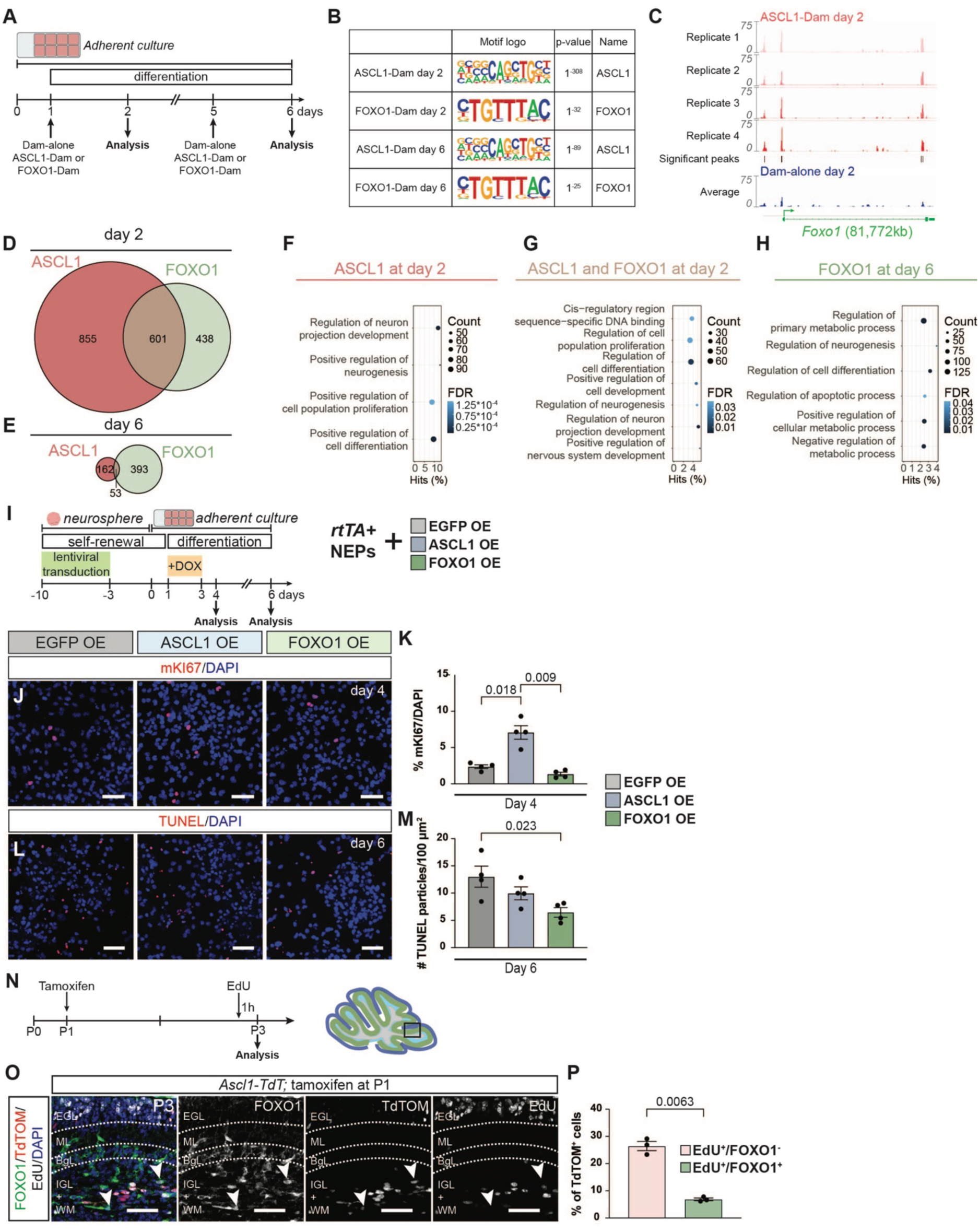
ASCL1 and FOXO1 independently regulate proliferation and cell survival during *in vitro* NEP differentiation. (A) Experimental design. (B) Table of HOMER results showing ASCL1 and FOXO1 binding motifs and significance in respective conditions (Table S3). (C) ASCL1-Dam tracks and the average of Dam-alone track (n=4) on day 2 of differentiation at the *Foxo1* loci. Significant ASCL1-Dam peaks are shown (FDR < 0.05). (D-E) Venn diagrams of significant and reproducible peaks comparing ASCL1-Dam and FOXO1-Dam on day 2 (D) and 6 (E) of differentiation (Table S4). (F-H) Selected significantly enriched GO terms for genes with ASCL1-Dam binding at day 2 (F), ASCL1-Dam and FOXO1-Dam binding at day 2 (G) or FOXO1-Dam binding at day 6 (H) are shown (Table S4). The circle size indicates the number of genes from a given GO term that contains a significant peak. The x-axis is the percentage of all the genes within a GO term with a significant peak. The colour gradient indicates the false discovery rate. (I) Experimental design. (J-K) Immunofluorescent analysis and quantification of EGFP OE, ASCL1 OE and FOXO1 OE cultures at day4 for proliferation marker mKi67 (one-way ANOVA, F_(1, 3)_=46.00, p=0.0029, n=4). (L-M) TUNEL staining and quantification of TUNEL^+^ particles on EGFP OE, ASCL1-OE and FOXO1-OE cultures at day 6 (one-way ANOVA, F_(2, 9)_=5.475, p=0.0278, n=4). (N) Experimental design. (O) Immunofluorescent analysis of P3 *Ascl1-TdT* brains that were given Tamoxifen at P1 and EdU one hour before the experimental endpoint. Arrowheads show TdTOM^+^/FOXO1^+^/EdU^-^ cells. (P) Quantification of EdU^+^ and EdU^+^/nuclear FOXO1^+^ cells in all TdTOM^+^ cells at P3 (n=3, paired Student’s t-test). Representative images are shown. Graphs show mean ± s.e.m. and significant p-values for multiple comparisons are shown in the figure. Scale bars: 50 µm.

### ASCL1 and FOXO1 have shared and distinct binding targets during differentiation

Having established that ASCL1 and FOXO1 are important for NEP-to-inhibitory neuron differentiation, we took a targeted DNA adenine methyltransferase identification (Targeted DamID; TaDa) approach (Marshall and Brand, 2015; Marshall et al., 2016; Southall et al., 2013) to identify their direct regulatory targets. Primary NEPs were infected with lentiviral constructs for expression of untethered Dam or fusions of either ASCL1 or FOXO1 with Dam one day before analysis on day 2 and 6 of differentiation (Fig. 4A). Time points were decided based on peak ASCL1 and FOXO1 expression during differentiation (Fig. 2K-L). After normalisation of fusion protein data to their respective Dam-alone control, significant peaks were calculated. Motif enrichment analysis revealed a significant enrichment of the ASCL1 and FOXO1 motifs within significant peaks derived from their respective binding data at both timepoints, confirming the faithful binding of the fusion proteins (Fig. 4B and Table S3).

ASCL1 OE NEPs showed a significant upregulation of FOXO1 during *in vitro* differentiation (Fig. 3G), suggesting that FOXO1 may be a direct transcriptional target of ASCL1. Therefore, we assessed the *Foxo1* locus for ASCL1-Dam binding. On day 2, ASCL1 binding was present at the FOXO1 promoter, indicating direct regulation of FOXO1 expression (Fig. 4C). To further resolve the targets of ASCL1 and FOXO1, we next identified all genes that showed significant ASCL1 or FOXO1 binding peaks within ±1kb of their transcriptional start site (Fig. 4D-E). We observed that more genes were bound by ASCL1 and/or FOXO1 on day 2 compared to day 6 (ASCL1: 1456 vs. 215 genes, FOXO1: 1039 vs. 446) (Fig. 4D-E). Interestingly, on day 2 of differentiation, a greater proportion of genes were associated with significant ASCL1 binding (1.40-fold compared to FOXO1), whereas this was inverted on day 6, with FOXO1 being associated with more genes (2.07-fold compared to ASCL1) (Fig. 4D-E). This suggests that ASCL1 has a more dominant role at early stages of differentiation, whereas FOXO1 has a more dominant role at later stages, in line with their temporal expression patterns we identified *in vitro* (Fig. 2K-L). Importantly, we observed a 31.73% and 8.72% overlap between genes bound by both ASCL1 and FOXO1 on days 2 and 6 of differentiation, respectively (Fig. 4D-E). This highlights potential cooperation between ASCL1 and FOXO1 during differentiation, though whether this is synergistic or antagonistic remains to be determined.

To provide further insights into the functional roles of genes bound by ASCL1 and FOXO1, we performed gene ontology analysis using Panther (Thomas et al., 2022) (Fig. 4F-H). Due to the low number of genes bound by ASCL1 and both ASCL1/FOXO1 on day 6, there were no significantly enriched GO terms identified (Table S4). Given the known role of ASCL1 during neurogenesis, it was unsurprising that genes bound by ASCL1 on day 2 were linked to GO terms related to regulation of nervous system development, neurogenesis and proliferation (Fig. 4F). These terms were also enriched for genes bound by both ASCL1 and FOXO1 at day 2 (Fig. 4G). In contrast, genes bound by FOXO1 only were enriched in GO terms related to metabolic and apoptotic processes (Fig. 4H, Table S4). These findings are in agreement with prior studies showing that FOXO1 regulates metabolic reprogramming, cell survival and apoptosis in other cell types (Bastie et al., 2005; Brunet et al., 2004; De La Torre-Ubieta et al., 2010; Puigserver et al., 2003; Schäffner et al., 2018; Yuan et al., 2008). Collectively, this data outlines key overlapping and independent roles of ASCL1 and FOXO1 during inhibitory neurogenesis. During the early stages of differentiation, FOXO1 cooperates with ASCL1 by targeting neurogenic genes, but later diverges in binding to target genes involved in survival and maturation.

### ASCL1 and FOXO1 regulate distinct cellular processes during differentiation

Our analysis suggests that ASCL1 and FOXO1 cooperate during the differentiation of NEPs to inhibitory neurons, regulating different cellular processes to facilitate inhibitory neuron production. Based on the GO term analysis of ASCL1 and FOXO1 bound genes, we hypothesised that alongside their roles in neurogenesis, ASCL1 may regulate proliferation and FOXO1 may regulate cell survival. To test this hypothesis, we assessed proliferation and apoptosis in ASCL1 OE and FOXO1 OE conditions compared to the EGFP OE control (Fig. 4I). While ASCL1 OE showed a significant increase in the percentage of mKI67^+^ cells at day 4 of differentiation (2.99 ± 0.40-fold compared to EGFP OE, n=4), FOXO1 OE did not affect proliferation (Fig. 4J-K). On the otherhand, FOXO1 OE cells showed a significant decrease in TUNEL^+^ particles compared to the EGFP OE control (0.50 ± 0.07-fold, n=4), while ASCL1 OE had no effect (Fig. 4L-M). This data suggests that the differentiation of NEPs to inhibitory neurons occurs via a proliferative neurogenic ASCL1^+^ state that transitions into a FOXO1^+^ state which then promotes neuronal survival and maturation. Indeed, analysis of proliferation in P3 *Ascl1-TdT* cerebella that were given tamoxifen at P1 revealed that, of the TdTOM^+^ cells, FOXO1^+^ cells showed a significant 0.26 ± 0.018-fold (n=3) reduction in EdU incorporation compared to FOXO1^-^ cells (Fig. 4O-P). This analysis confirmed the change in the proliferative status of NEPs from an ASCL1^+^ to a FOXO1^+^ state during differentiation. In summary, these results demonstrate that during NEP-to-inhibitory neuron differentiation, ASCL1 promotes proliferation, and FOXO1 supports cell survival, also explaining why ASCL1-OE promoted more neurogenesis than FOXO1-OE (Fig. 3J).

### Activation of WNT signalling promotes a FOXO1^+^ state and increases neurogenesis in primary NEPs

It remained unclear what promotes the transition of *Ascl1*-NEPs from the ASCL1^+^ state to a FOXO1^+^ state during differentiation. To explore this, we revisited our scRNA-seq data to identify signalling pathways with activation patterns overlapping with *Foxo1* expression and regulon activity (Figs 1H and 1K). To this end, we utilised Cell2Fate (Aivazidis et al., 2025), an algorithm optimised to reconstruct cellular trajectories from scRNA-seq data, which orders cells according to an inferred time and computes modules of genes that follow the same mRNA splicing dynamics (Figs 5A-B and S3A). This allowed us to interrogate commonalities between the genes driving the computed inferred time. The state of each module is classified for individual cells based on how the amount of spliced UMIs for genes within a module change over inferred time. Nine modules with distinct temporal dynamics were identified (Figs 5C-E, S3B-C and Table S5). Of those modules, we specifically focused on modules 2, 4 and 6 that are transiently active in the bipotent state (cluster W4), around the *Foxo1*^+^ state (cluster W3) or immediately following the *Foxo1*^+^ state (clusters W2 and W0) (Fig. 5C-E). GO term analysis of the top 200 module markers revealed an upregulation of WNT signalling prior to the *Foxo1*^+^ state (module 2) (Fig. 5F). This was followed by an enrichment of neurogenesis-related genes (module 4) and then genes involved in negative regulation of WNT signalling and cell migration/motility (module 6) (Fig. 5G-H and Table S5). This data suggests that transient activation of the WNT signalling pathway may facilitate the progression of NEP-to-inhibitory neuron differentiation during postnatal development.

**Fig. 5.**
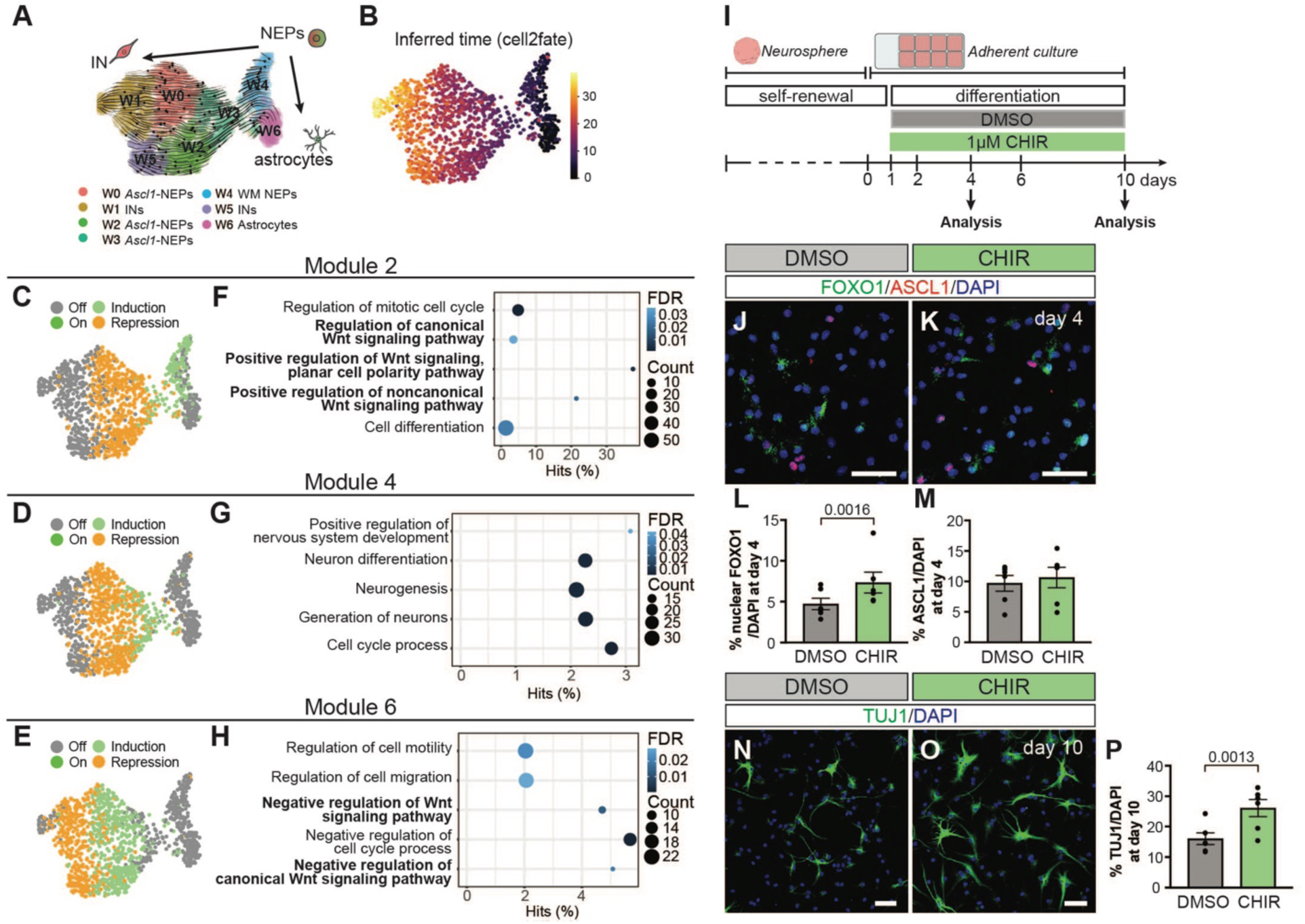
WNT signalling is a positive regulator of FOXO1 and neural production during NEP-to-inhibitory neuron differentiation. (A-B) UMAP projection of cellular trajectories (A) or inferred time (B) computed using Cell2Fate within the WM NEPs. (C-E) UMAP projections showing the transcriptional state of module gene for modules 2 (C), 4 (D) or 6 (E). (F-H) GO term analysis of module genes for modules 2 (F), 4 (G) or 6 (H). Selected significantly enriched GO terms are shown (Table S5). The circle size indicates the number of genes from a given GO term. The x-axis is the percentage of all the genes within a GO term. The colour gradient indicates the false discovery rate. (I) Experimental design. (J-M) Immunofluorescent analysis (J, K) and quantification of nuclear FOXO1^+^ (L) and ASCL1^+^ (M) cells in cultures treated with DMSO or CHIR at day 4 of differentiation (n=6, paired Student’s t-test). (N-P) Immunofluorescent analysis and quantification of TUJ1^+^ cells in cultures treated with DMSO or CHIR at day 10 of differentiation (n=6, paired Student’s t-test). Representative images are shown. Graphs show mean ± s.e.m. Scale bars: 50 µm.

To test whether WNT signalling promotes the transition of *Ascl1*-NEPs into the *Foxo1^+^* state, we activated canonical WNT signalling during *in vitro* differentiation with CHIR99021 (CHIR) (Fig. 5I). CHIR treatment during differentiation led to a significant 1.55 ± 0.27-fold (n=6) increase in the percentage of cells with nuclear FOXO1 at day 4 but did not significantly affect the number of ASCL1^+^ or mKI67^+^ cells (Figs 5J-M and S3D). CHIR treatment also resulted in a significant 1.62 ± 0.17-fold (n=6) increase in the percentage of TUJ1^+^ cells produced after 10 days of differentiation (Fig. 5N-P). Collectively, this data suggests that WNT signalling promotes the transition from the ASCL1^+^ state to the FOXO1^+^ state during *Ascl1*-NEP differentiation to inhibitory neurons during postnatal cerebellar development.

### The ASCL1-FOXO1 axis is conserved during human NEP-to-inhibitory neuron differentiation *in vitro*

To assess whether the molecular mechanisms that promote mouse molecular layer inhibitory neuron differentiation are conserved in the human cerebellum, we utilised human fetal cerebellar tissue to study the expression of ASCL1 and FOXO1. Immunofluorescent analysis on histological sections from a 17 postconception week (pcw, an equivalent stage to around birth in mice (Haldipur et al., 2022)) human cerebellum revealed nuclear FOXO1 expression in SOX2^+^ cells in the prospective WM, some of which were also ASCL1^+^. This suggested that FOXO1 is expressed in human *ASCL1*-NEPs (hNEPs) (Fig. 6A).

**Fig. 6.**
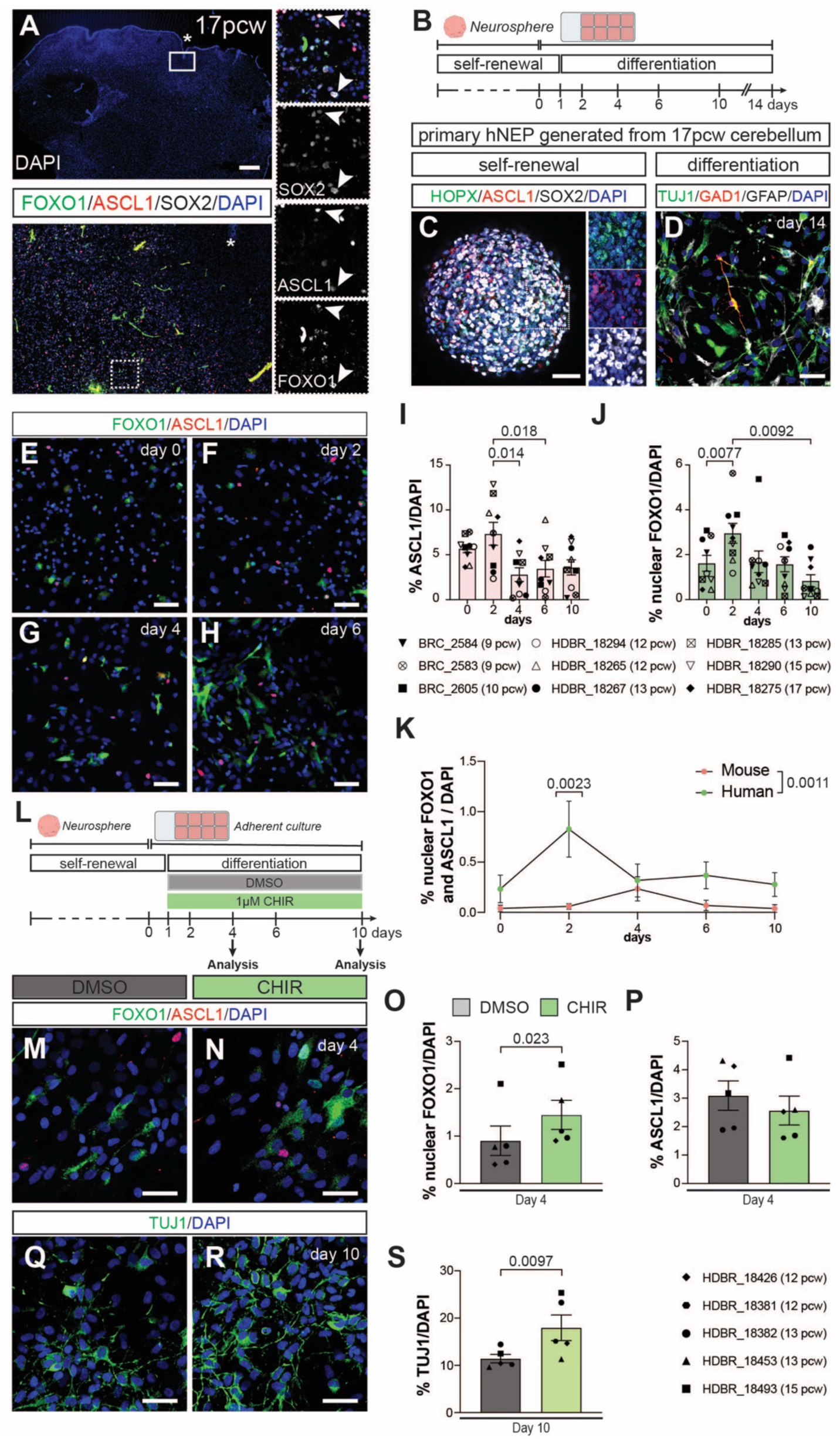
The role of WNT signalling in promoting neural production via FOXO1 is conserved in human cerebellar NEP differentiation. (A) Immunofluorescent analysis of a 17 post-conception week (pcw) human cerebellum. Arrowheads show FOXO1^+^/ASCL1^+^/SOX2^+^ triple positive cells; asterisks indicate an anchoring centre. (B) Experimental design. (C-D) Immunofluorescent analysis of primary human NEP (hNEP) neurospheres showing SOX2^+^, HOPX^+^ and ASCL1^+^ cells (C) and 2D cultures on day 14 of differentiation showing GFAP^+^, TUJ1^+^ and GAD1^+^ cells (D). Insets show high magnification images. (E-H) Immunofluorescent analysis for ASCL1 and FOXO1 during hNEP differentiation. Representative images of day 0 (H), 2 (I), 4 (J) and 10 (K) are shown. (I-J) Quantification of ASCL1^+^ (I) and nuclear FOXO1^+^ (J) cells throughout *in vitro* differentiation (one-way ANOVA, I: F_(2, 23)_=6.614, p=0.0023, n=9. J: F_(2, 20)_=6.812, p=0.0033, n=9). (K) Quantification of nuclear FOXO1^+^ and ASCL1^+^ double positive cells during *in vitro* differentiation of mouse and human NEPs (two-way ANOVA, I: F_(1, 70)_=11.51, p=0.0011, n=7 (mouse) or 9 (human). (L) Experimental design. (M-P) Immunofluorescent analysis and quantification of nuclear FOXO1^+^ (O) and ASCL1^+^ (P) cells in hNEP cultures treated with DMSO or CHIR at day 4 of differentiation (n=5, paired Student’s t-test). (Q-S) Immunofluorescent analysis and quantification of TUJ1^+^ cells in hNEP cultures treated with DMSO or CHIR at day 10 of differentiation (n=5, paired Student’s t-test). Representative images are shown. Graphs show mean ± s.e.m. and significant p-values for multiple comparisons are shown in the figure. Scale bars: 50 µM, except for A (500 µM).

To investigate the dynamics of ASCL1 and FOXO1 expression during hNEP differentiation, we established primary NEP cultures from human fetal tissue ranging from 9-17 pcw. Similar to primary mouse NEPs, these cells were also maintained as neurospheres with EGF and FGF2 and were differentiated as adherent cultures by growth factor withdrawal and 2% FBS treatment (Fig. 6B). Immunofluorescent staining of hNEP neurospheres revealed that most cells expressed SOX2, and both HOPX^+^ and ASCL1*^+^*hNEPs were present (Fig. 6C). Following differentiation for 14 days, we obtained a mixed culture of GFAP^+^ astrocytes and TUJ1^+^ neurons, some of which were GAD1^+^ inhibitory neurons (Fig. 6D). This data confirms that hNEPs can also generate inhibitory neurons *in vitro,* providing us with a reliable model to study differentiation mechanisms. We then assessed the percentage of ASCL1^+^ and nuclear FOXO1^+^ cells during *in vitro* differentiation (Fig. 6E-J). We observed that the percentages of both ASCL1^+^ and nuclear FOXO1^+^ cells increase early during differentiation (day 2) and then reduce to significantly lower levels (Fig. 6I-J). At the beginning of differentiation, the percentages of ASCL1^+^ and nuclear FOXO1^+^ cells were higher than at the same stage during mouse NEP differentiation (Fig. 6I-J, compared to Fig. 2K-L). Interestingly, the highest expression of ASCL1 and FOXO1 overlapped at day 2, unlike what was observed during mouse NEP differentiation. In line with this observation, we also detected a 13.76 ± 4.83-fold (n=9) increase in the percentage of ASCL1^+^/FOXO1^+^ double-positive cells on day 2 of hNEP differentiation compared to mouse NEPs (Fig. 6K). In summary, while ASCL1 and FOXO1 are transiently upregulated during differentiation of hNEPs, the transient ASCL1^+^/FOXO1^+^ double positive state seems to be prolonged in hNEP differentiation *in vitro* compared to what is observed in mice.

Finally, to test whether WNT signalling promotes ASCL1^+^ to FOXO1^+^ transition during *in vitro* hNEP differentiation, we treated the differentiating hNEPs with 1µM CHIR (Fig. 6L). Activation of WNT signalling led to a significant 1.60 ± 0.34-fold (n=5) increase in nuclear FOXO1^+^ at day 4 of differentiation, while the percentage of ASCL1^+^ cells did not change (Fig. 6M-P), consistent with our observations in mouse NEPs. Importantly, activation of WNT signalling during *in vitro* differentiation of hNEPs also increased the percentage of TUJ1^+^ cells at day 10 of differentiation 1.57 ± 0.24-fold (n=5) compared to the DMSO controls (Fig. 6Q-S). Collectively, this data suggests that the ASCL1-WNT-FOXO1 axis is conserved during NEP-to-inhibitory neuron differentiation between mice and humans, while the dynamics of the transient FOXO1^+^ state may differ between the species.

## Discussion

Resolving differentiation programmes is crucial for understanding how the nervous system is built in a region-specific manner. This is particularly important where imbalances in one neural type could lead to devastating defects in brain physiology. For example, the molecular layer inhibitory neurons of the cerebellum play a crucial role in modulating Purkinje neuron firing, and hence, cerebellar function (Brown et al., 2019). In addition, molecular layer inhibitory neuron numbers are affected in cerebellar disorders such as ataxias (Edamakanti et al., 2018). However, the molecular mechanisms that regulate cerebellar molecular layer inhibitory neuron differentiation remain poorly understood. In this study, we established novel aspects of the GRN that promotes NEP-to-inhibitory neuron differentiation during postnatal cerebellar development. We used analysis of scRNA-seq from NEPs isolated from the postnatal mouse cerebellum to identify core genes in the GRNs that promote gliogenic and neurogenic differentiation of WM NEPs. We found that WM *Ascl1*-NEPs utilise a combinatorial transcriptional network and resolved the context-dependent functions for ASCL1 and FOXO1, two key neurogenic transcription factors. Our transcriptional and targeted DamID data suggest that both ASCL1 and FOXO1 promote neurogenesis with temporally dynamic, shared and divergent functions. Both transcription factors facilitate a neurogenic programme, whilst ASCL1 independently regulates proliferation and FOXO1 expression. FOXO1, on the other hand, promotes survival during NEP differentiation separately from ASCL1. Our data also suggests that WNT signalling induces NEP-to-inhibitory neuron differentiation by facilitating the transition of the *Ascl1*-NEPs from an ASCL1^+^ state to a transient FOXO1^+^ cell state. Using primary hNEP cultures generated from fetal human cerebella, we then demonstrated that the ASCL1-FOXO1 differentiation axis and the role of WNT signalling in hNEPs-to-inhibitory neuron differentiation is conserved. Based on this work, we propose a model for the role of FOXO1 and its interaction with ASCL1 during NEP-to-inhibitory neuron differentiation in the developing cerebellum (Fig. 7). This work provides unique insights into how cerebellar inhibitory neuron production is regulated while resolving context-dependent functions of ASCL1 and FOXO1 in mouse and human NEPs.

**Fig. 7.**
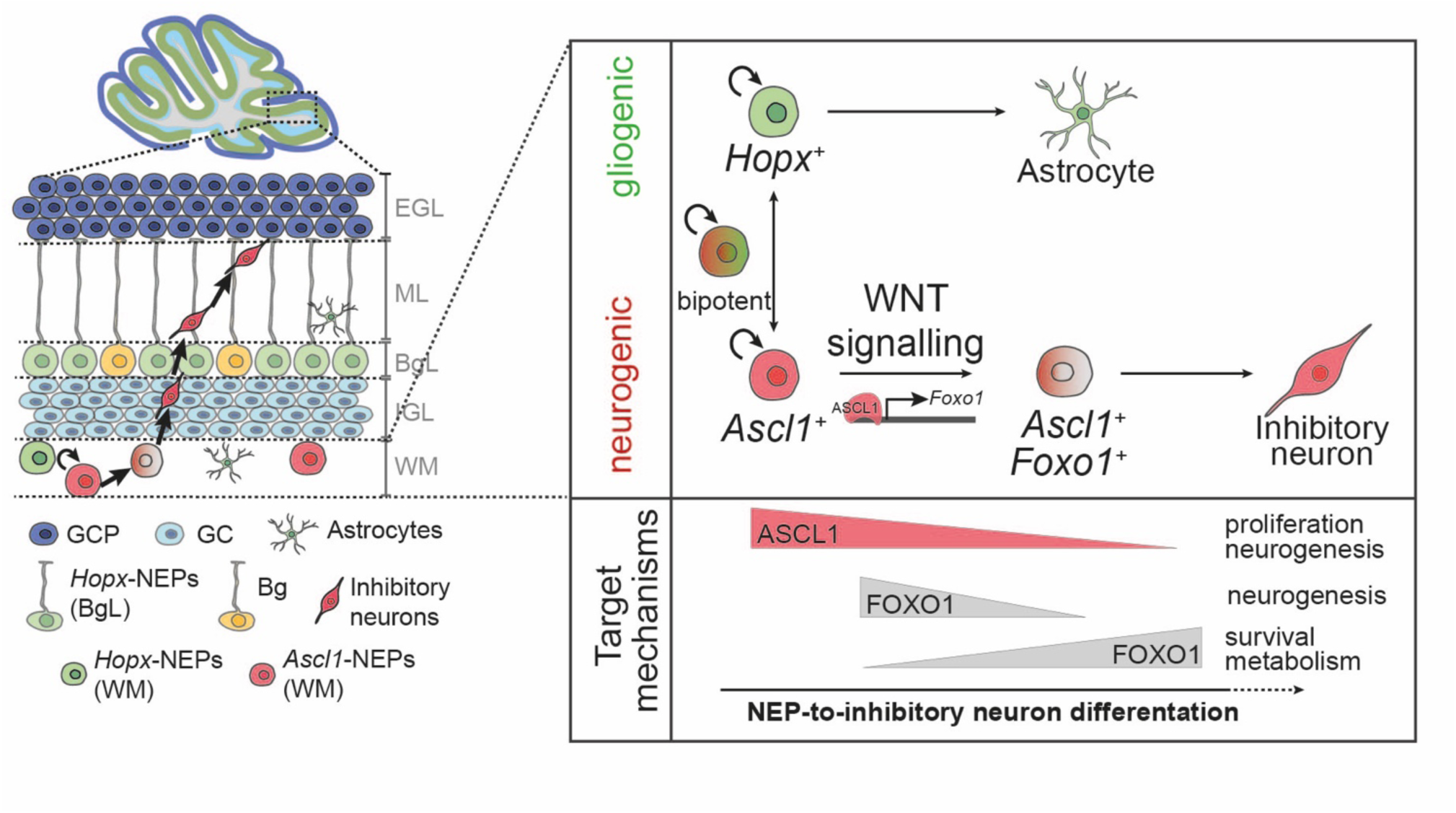
Summary. Working model for how *Ascl1*-NEPs transition to a FOXO1^+^ state via WNT signalling activation to drive NEP-to-inhibitory neuron production in the cerebellum and the downstream targets of ASCL1 and FOXO1 during cerebellar molecular layer inhibitory neuron differentiation provides insights into their context-dependent functions.

The function of transcription factors is highly context-dependent, which has allowed them to be evolutionarily reused across different cell types and developmental stages (Argelaguet et al., 2022). Using lentiviral overexpression, we demonstrated that sustained ASCL1 and FOXO1 overexpression in primary NEPs increases neuron production *in vitro* (Fig. 3J). The sustained overexpression may have disrupted some of the GRNs driving NEP differentiation and could lead to impaired differentiation and maturation. Additionally, our lentiviral overexpression approach is not cell type-specific, resulting in ASCL1 and FOXO1 overexpression in different NEP subtypes *in vitro*. Therefore, some of the observed differences between ASCL1 and FOXO1 overexpression, such as the differences in the morphology of TUJ1^+^ cells could be attributed to these technical challenges. *In silico* GRN reconstruction of the scRNA-seq data suggested a low level of FOXO1 regulon activity in bipotent NEPs and astrocytes. The role of FOXO1 in these cell types remains to be determined. FOXO1 has been reported to regulate cellular metabolism and inhibit ROS generation in astrocytes in other brain regions (Doan et al., 2023; Wang et al., 2022). Thus, we can speculate that the function of FOXO1 may partially be conserved between cerebellar interneurons and astrocytes.

Our data shows that ASCL1 and FOXO1 bind to a shared group of genes related to neurogenesis during *in vitro* NEP-to-inhibitory neuron differentiation while independently regulating proliferation and cell survival, respectively, at different stages. These observations are consistent with the function of ASCL1 and FOXO1 in other cell types and, additionally, other FOXO family transcription factors were shown to functionally interact with ASCL1 at the same target genes (Bastie et al., 2005; Brunet et al., 2004; Castro et al., 2006; De La Torre-Ubieta et al., 2010; Imayoshi et al., 2013; Park et al., 2017; Webb et al., 2013). However, it remains unclear whether ASCL1 and FOXO1 are synergistic and/or antagonistic at loci they both bind.

Cell2Fate analysis revealed dynamic regulation of WNT signalling during NEP-to-inhibitory neuron differentiation (Fig. 5C-H). We demonstrated that CHIR treatment during *in vitro* mouse and human NEP differentiation increases nuclear FOXO1^+^ cells and neuron production while not affecting the frequency of ASCL1^+^ cells (Figs 5J-P, 6M-S). Similarly, overexpression of ASCL1 during *in vitro* differentiation resulted in an increased amount of nuclear FOXO1^+^ cells and neuron production (Fig. 3F-G). The mechanisms of how ASCL1 and WNT signalling increase nuclear FOXO1^+^ levels converge remain to be investigated. ASCL1 has been shown to promote WNT signalling by directly repressing DKK1, a negative regulator of WNT signalling, and synergising with WNT3A to induce active WNT signalling in glioblastoma stem cells (Rheinbay et al., 2013). Thus, ASCL1 may prime *Ascl1*-NEPs for WNT signalling. In other systems, WNT signalling has been shown to both positively and negatively regulate FOXO1 in a context-dependent manner (Cheng et al., 2024; Okada et al., 2015; Sreekumar et al., 2017). In a conditional knockout mouse using *Nes-Cre* to delete WNT5a, a non-canonical WNT ligand, the production of molecular layer interneurons in the postnatal cerebellum was decreased (Subashini et al., 2017). How the canonical and noncanonical WNT signalling crosstalk promotes NEP-to-inhibitory interneuron differentiation in the postnatal cerebellum remains to be determined. Interestingly, activation of WNT signalling in cerebellar progenitors during embryonic development causes abnormal cerebellar development and a reduced number of inhibitory neurons (Pei et al., 2012; Yang et al., 2019). This highlights the multifaceted role of WNT signalling in cerebellar development, with possibly distinct roles in embryonic and postnatal developmental stages. Furthermore, a down-regulation of WNT-related genes was observed after the *Foxo1*^+^ state in the WM NEP transcriptomes (Fig. 5E-H). FOXO1 has been reported to suppress WNT signalling in neural stem cells(Paik et al., 2009) and other cell types (Guan et al., 2015; Iyer et al., 2013). It is plausible that WNT signalling and FOXO1 form a negative feedback loop as part of the postnatal NEP-to-inhibitory neuron differentiation program.

Finally, to establish to what extent these mechanisms are conserved in human cerebellum development, we adapted our *in vitro* differentiation paradigm to establish primary hNEP cultures from fetal cerebella. We found that the ASCL1-FOXO1 axis and the role of WNT signalling are conserved in hNEP-to-inhibitory neuron differentiation (Fig. 6E-S). Interestingly, we did observe differences in the temporal dynamics of ASCL1 and FOXO1 compared to the mouse NEP differentiation, where the proportion of ASCL1^+^/FOXO1^+^ double-positive cells was significantly higher during hNEP differentiation (Fig. 6K). We speculate that the prolonged double-positive state may be an evolutionary adaptation to scale up the production of molecular layer interneurons to support the expanded human cerebellar hemispheres (Kebschull et al., 2020). Our analysis was performed on samples obtained from a wide range of ages (9-17 pcw). We did not find any correlation with the age of the donor. However, it remains unclear whether different cell states or cell types are captured at different ages and how this might influence *in vitro* differentiation. Advancements in human pluripotent stem cell-derived cerebellar organoids may offer an alternative platform to functionally test the molecular mechanisms driving hNEP differentiation (Atamian et al., 2024; Brás et al., 2022). Collectively, our work uncovers parts of the GRN driving NEP-to-molecular layer inhibitory differentiation in mice and humans during late cerebellar development. These findings provide insights into how neurogenesis is regulated in a neuron type- and region-specific manner by resolving the context-dependent functions of ASCL1 and FOXO1. Our results have the potential to inform future therapies for disorders where inhibitory neuron production is impaired.

### Limitations

While our *in vitro* use of primary cells has proven a powerful tool to identify molecular mechanisms of NEP behaviour in mice and humans, these findings would need to be further tested using conditional knockout models for FOXO1. Importantly, the low number of relevant cells (*Ascl1*-NEPs) that can be isolated from the neonatal mouse brain impairs molecular studies on cells freshly isolated from the brain, further highlighting the utility of our *in vitro* models. Finally, the limited availability of human fetal tissue, particularly from later stages, and the inability to maintain cultures long-term pose a significant challenge to characterising developmental dynamics and performing functional validation in hNEPs.

## Methods

### Animal husbandry

All the mouse experiments were performed according to the protocols approved by Home Office Project Licence PP8035780. The mouse lines used in this study are: *Nes-CFP* (Mignone et al., 2004; Wojcinski et al., 2017), *Ascl1^CreERT2^* (Pacary et al., 2011; Sudarov et al., 2011) and *Rosa26^lox-stop-lox-TdTomato^* (ai14, stock no. 007909, The Jackson Laboratory) (Takeda et al., 2011). Animals were maintained on an outbred *Swiss Webster* or *CD1* background, housed on a 12-hour light/dark cycle and had access to food and water ad libitum. Both sexes were used for the study.

For genetic inducible fate mapping, P1 pups were injected subcutaneously with 200 µg/g Tamoxifen (Sigma-Aldrich, T5648). For proliferation analysis, 50 µg/g EdU (ThermoFisher, E10187) was injected subcutaneously one hour prior to the experimental endpoint.

### Human fetal tissue

9-17 pcw old human fetal cerebellar tissue was obtained from either the Cambridge University Hospitals NHS Foundation Trust under permission from NHS Research Ethical Committee (96/085) or the MRC/Wellcome Trust Human Developmental Biology Resource (London (University College London (UCL) site REC reference: 18/LO/0822) and Newcastle (Newcastle site REC reference: 18/NE/0290), Project no: 200702 (www.hdbr.org). Only samples with no genetic abnormalities were used.

### Primary NEP neurosphere cultures

Cerebella from *Nes-Cfp/+* or *Swiss Webster* pups were dissected and dissociated using Accutase (ThermoFisher, #A1110501) for 10 minutes at 37_°_C. Once a single cell suspension was obtained, cells were washed with NSC media (Neurobasal (ThermoFisher, #10888022) with 1X B27 (ThermoFisher, #A3353501), 1X N2 (ThermoFisher, #11520536), 2mM L-glutamine (ThermoFisher, #A2916801), 1X non-essential amino acids (ThermoFisher, 11140035) and 100 U/mL penicillin-streptomycin (ThermoFisher, #15140122)) and plated onto ultra-low attachment plates ∼100,000 cells/mL density. Cultures were maintained with 20 ng/mL of EGF (Fisher Scientific, #PMG8041) and FGF2 (Qkine, #Qk042-0500).

Primary hNEP neurospheres were established similarly to the mouse NEPs, except for an additional filtering step with a 40 µM nylon filter (Merck, #CLS352340) after dissociation to remove debris. hNEP cultures were maintained in the NSC media supplemented with human EGF (QKINE, #Qk011-0100) and FGF2 (QKINE, #Qk027-0100). Growth factors were supplemented every other day and media was changed weekly.

### *In vitro* differentiation

Chamber slides (Thistle Scientific, #IB-80806) were coated with poly-D-lysine and laminin. Briefly, 8-well slides were incubated with 250 µL poly-D-lysin (ThermoFisher, #A389040) for >2 hours at 37_°_C. The plates were rinsed with PBS+/+ (+Mg and +Ca, ThermoFisher, #14040091) twice and then were incubated with 2 µg/cm^2^ laminin (Merck, #L2020) or human laminin-521 (Thermo Fisher, #A29249) in PBS+/+ (ThermoFisher, #14040091). The plates were stored at 4_°_C until use. On the day of differentiation, the plates were incubated for >1 hour at 37_°_C and washed once with PBS+/+ prior to cell seeding.

Neurospheres were dissociated using accutase and washed with NSC media once. Cells were plated at a density of 250,000 cells/cm^2^ (mouse) or 100,000 cells/cm^2^ (human) on poly-D-lysine and laminin-coated plates. Cells were kept in self-renewing conditions (NSC media + growth factors) overnight and then switched to differentiation media (NSC media + 2% fetal bovine serum (FBS, ThermoFisher, #10438026)). The media was changed every other day and on days prior to fixing the cells. For WNT activation, cells were treated with 1µM CHIR99021 (BioTechne, #4423/10). 0.1% DMSO was used as a control. HDBR_18426 (12 pcw), HDBR_18381 (12 pcw), HDBR_18453 (13 pcw), HDBR_18382 (13 pcw) and HDBR_18493 (15 pcw) were used in the human WNT experiments. 2-3 technical replicates were performed for each sample and averaged.

Cells were fixed at room temperature (15 minutes for 2D cultures and 30 minutes for neurospheres using 4% paraformaldehyde (PFA, ThermoFisher Scientific, #043368.9M) and washed thrice with 1X PBS. Fixed cells were stored at 4_°_C until immune fluorescent analysis.

### Tissue preparation for histological analysis

The dissected neonatal mice brains or fetal human cerebellum were drop fixed in 4% PFA at 4_°_C for up to a week followed by 30% sucrose (Merck, #1076511000) in PBS until the brains sunk. Then, the brains were embedded in OCT (VWR, #361603E) blocks and stored at −80_°_C until further processing. 14µm thick sections were obtained using a cryostat (Leica CM3050S) and stored at −80_°_C.

### Immunofluorescent analysis

Tissue sections, neurospheres and 2D cultures were subjected to immunofluorescent analysis. The list of antibodies used for this analysis is in Table S6. Slides were allowed to come to room temperature. Samples were blocked with blocking buffer (5% Bovine serum albumin (Merck Life Science, A9418-50G) in 1X PBS and 0.1% triton-X). For ASCL1 staining on slides, mouse-on-mouse blocking was performed per the manufacturer’s guidelines (ThermoFisher, #R37621). Primary antibody incubation was performed in blocking buffer overnight at 4_°_C. Samples were then washed three times with 1X PBS with 0.1% triton-X for 5 minutes. Secondary antibody incubation was performed in blocking buffer for 1-2 hours at room temperature, and samples were protected from light for the rest of the procedure. After secondary antibody staining cells were washed three times with 1X PBS with 0.1% triton-X for 5 minutes and DAPI (Sigma-Aldrich, MBD0015-1ML) was used for counterstain. Sections were mounted with fluoro-gel (EMS, #17985-10). Cultures were stored in 1X PBS. Cells and sections were stored at 4_°_C in the dark until imaging.

A similar procedure was used for neurosphere staining with minor modifications to the blocking buffer (PBS containing 0.2% bovine serum albumin, 0.02% Sodium Dodecyl Sulfate (SDS) and 0.3% Triton-100X) and extended incubation times.

Apoptosis was detected using TUNEL staining per manufacturer’s guidelines (ThermoFisher, 16314015). Where relevant, EdU was detected using Click-it assay with sulfo-azide-Cy5 (Lumiprobe Corporation, A3330).

### Image acquisition and analysis

Images were acquired using either a Leica CKX53 epifluorescence microscope, Zeiss LSM880 confocal or Andor BC43 CF confocal microscope. Images were processed using ImageJ. Images within an experiment were acquired with identical settings and max intensity projections are shown in the figures.

Quantification of tissue sections was performed on lobules 4 and 5 in three midsagittal sections/brain. Quantification was done manually using ImageJ/FIJI Cell Counter Plugin (NIH). For experiments that involved 2D cultures, three images were acquired from each well and 3-10 wells/condition were quantified across >3 individual experiments. Quantification was done manually using ImageJ/FIJI Cell Counter Plugin (NIH). TUNEL quantification was done with FIJI, converting the images to an 8-bit image, and quantifying particles with the Analyze Particles tool.

### Lentiviral production

VSV-G pseudotyped lentivirus was produced per standard protocols using HEK293T cells and the packaging plasmids for Gag/pol, VSV-G, REV and Tat plasmids (ratio 1:2:1:1) were added together with 5 µg of transfer plasmid. Upon collection, the media was filtered through a 0.2 µm filter (Merck, #SLGSVR255F) and concentrated using Lenti-X reagents (Takara, #631232) per manufacturer’s guidelines. The concentrated lentivirus was resuspended a 300 µL of Opti-MEM (ThermoFisher, #11058021) per 10 cm dish. The virus was aliquoted in appropriate volumes, snap-frozen on dry ice and stored at −80_°_C until use. Viral titers were calculated using ELISA. The following transfer vectors were used for this study:

| Transfer vector | Reference | Catalogue number |
| --- | --- | --- |
| Tet-O-FUW-Ascl1 | Liu *et al* (Liu et al., 2018) | Addgene cat# 27150 |
| Tet-O-FUW-Foxo1 | Liu *et al* (Liu et al., 2018) | Addgene cat# 118588 |
| FUW-tetO-EGFP | Panciera *et al* (Panciera et al., 2016) | Addgene cat# 84041 |
| Lenti-SFFV-mNG-DamAlone | Lim *et al* (Lim et al., 2023) | N/A |
| Lenti-SFFV-mNG-Dam-Ascl1 | This paper | N/A |
| Lenti-SFFV-mNG-Foxo1-Dam | This paper | N/A |
| pLV-EF1a-IRES-Puro | Hayer *et al* (Hayer et al., 2016) | Addgene cat# 85132 |
| Scramble shRNA | Sarbassov *et al* (Sarbassov et al., 2005) | Addgene cat# 1864 |
| pLKO.1 | Moffat *et al* (Moffat et al., 2006) | Addgene cat# 10878 |

### Lentiviral infection *in vitro*

Primary NEP neurospheres were dissociated into single cells as described above and incubated with the respective virus at 1-10 multiplicity of infection (MOI) and 4 µg/mL protamine sulfate (Merck, #1101230005) for 12-18 hours.

For the overexpression experiments, a stable rtTA NEPs cell line was first established and then reinfected with the subsequent overexpression virus prior to analysis. These cells were differentiated as described above 3-10 days after infection, with the exception that Tet system approved FBS (ThermoFisher Scientific, A4736401) was used in the differentiation media. In the doxycycline condition, 1 µg/mL doxycycline (Sigma, #D9891) was added to the differentiation media from days 1-3 (Fig. 3A).

For shRNA mediated knockdown, 4 different *Foxo1* shRNA (*Foxo1*-sh1: CCGCCAAACACCAGTCTAAAT, *Foxo1*-sh2 CGGAGGATTGAACCAGTATAA, *Foxo1*-sh3 TGTAATGATGGGCCCTAATTC, *Foxo1*-sh4 CATGGACAACAACAGTAAATT) and a scrambled shRNA oligo were cloned into the pLKO.1 vector per guidelines(Moffat et al., 2006). shRNAs were designed using the Broad Institute TRC shRNA design tool. NEPs were infected and selected with 1ug/mL puromycin for 1 week to establish stable cell lines. Following infection, *Foxo1* expression levels were assessed by qPCR or immunofluorescence and 2/4 shRNAs (*Foxo1*-shRNA1 and *Foxo1-*shRNA2) that showed better knockdown were carried on for further analysis (Fig. 3L).

### Targeted DamID

FOXO1 or ASCL1 CDS was inserted into the SFFV-mNeonGreen-Dam lentiviral vector (Lim et al., 2023) by Gibson assembly. Differentiations were performed in 6-well plates (ThermoScientific, #140675). Differentiating NEPs were infected with Dam-alone, ASCL1-DAM or FOXO1-Dam lentivirus in neural stem cell media with 2% fetal bovine serum and 4 µg/mL protamine sulfate and were incubated at 37_°_C for 6 hours. Following infection, the media was changed to differentiation media and cells were collected for downstream analysis after 24 hours (Fig. 4A). Briefly, cells were dissociated with Accutase, washed once, and pellets were collected and stored at −80_°_C until library preparation.

Genomic DNA was extracted from cell pellets and processed as previously described (Marshall et al., 2016). Sequencing was performed by the Gurdon Institute NGS core as single end 100 bp reads on an Illumina NovaSeq 6000. Raw fastq files were analysed with a modified version of the damidseq_pipeline (Marshall and Brand, 2015). Reads were mapped to mm10 genome assembly, indexed using Bowtie2 and binned into 5’-GATC-3’ fragments. Each fusion protein (FOXO1/ASCL1) sample was normalised against each replicate of Dam-alone treatment, then averaged using RPM normalisation, in 300 bp bins. Binding intensity values were quantile normalised across all replicates for each stage and backtransformed. Files were converted to bigwig format using bedGraphToBigWig (v4) and imported into the Integrative Genomics Viewer (IGV v2.17.4) for visualisation. Macs2 (Zhang et al., 2008) (v2.1.2) was used to call broad peaks for fusion and Dam-alone pairs using the bam file output of the damidseq_pipeline. A FDR-threshold of <0.05 was used to identify significant binding peaks. For downstream analyses, significant peaks were further filtered to those present only in 3/4 replicates (repropeaks) using bedtools (Quinlan and Hall, 2010) (v2.26.0).

The repropeak bed files were read into R using the ChIPpeakAnno (Zhu et al., 2010) (v3.32.0) function toGRanges with format = ‘BED’, header = T and otherwise default parameters. The Grange objects were annotated individually using the function annotatePeakInBatch with AnnotationData = genes (EnsDb.Mmusculus.v79). All peaks where distancetoFeature was ± 1000 were discarded, and features with multiple peaks were then made unique. A list of the genes with a significant peak ± 1kb of the transcriptional start site was made and used to compute unique and shared genes using ggven (v0.1.14) with default parameters. A conversion table was made of ENSEMBL IDs and gene symbols using BiomaRt (Smedley et al., 2009) (v2.54.1) getBM specifying attributes=c(’ensembl_gene_id’, ‘ensembl_transcript_id’, ‘mgi_symbol’) and mart = useMart(biomart=“ENSEMBL_MART_ENSEMBL”, dataset = “mmusculus_gene_ensembl”, host = “https://www.ensembl.org/”, verbose = F). The gene set was exported as a csv-file using write.csv with default parameters. GO term analysis was performed using PANTHER (Thomas et al., 2022) (v19.0) via the online GUI available at https://pantherdb.org/. GO term analysis for biological process, molecular function and cellular component terms was performed individually with default parameters using Fischer’s Exact test and a false discovery rate calculation to correct for multiple testing. For individual transcription factors, all genes with a reproducible significant peak ±1kb of the transcriptional start site were used for the analysis, irrespective of the other transcription factor binding. For the co-bound analysis, genes must have a reproducible significant peak ±1kb from the transcriptional start site for both transcription factors at the given timepoint.

To perform motif enrichment analysis, the produced bed-files were loaded using toGRanges and annotated as described above. From the Grange object, a HOMER-compatible data frame was constructed. The HOMER-compatible data frame was exported as a bed-file using write.table specifying quote=F, sep=“\t”, row.names=F, col.names=F. Motif enrichment analysis was then performed using the HOMER(Heinz et al., 2010) (v5.0.1) function findMotifsGenome.pl using the mm10 genome specifying -mask and otherwise default parameters. The reported results are from the knownResults output.

### Single-cell RNA sequencing and data analysis

#### 10X CellPlex Single-cell RNA-sequencing

Sample preparation, cell multiplexing and droplet preparation, and mapping and initial quality control were performed as described in Pakula *et al*., 2025 (Pakula et al., 2025).

#### Clustering

NonIR (control condition in Pakula *et al*., 2025) cells from P1-3 and 5 cerebella were subsetted from the rds object using Seurat (Hao et al., 2024) (v4.1). The object was then split into a list based on the individual biological replicates using SplitObject and normalised using ScaleData with default parameters. The list of rds object was prepared for integration using SelectIntegrationFeatures and PrepSCTIntegration using default parameters, and subsequently integrated using IntegrateData with default parameters. The integrated rds object was normalised using SCTransform with vars.to.regress = c(“percent.mt”, “CC.Difference”, “nFeature_RNA”, “nCount_RNA”). Dimension reduction was then performed using RunPCA using all features in the integrated assay and npcs=100 and RunUMAP with parameters dims = 1:40, seed.use = 888, repulsion.strength = 0.1 and min.dist = 0.5 in addition to default parameters. Clustering was performed using FindNeighbors with default parameters and dims=1:30 and FindClusters with default parameters and a resolution varying between 0.1 and 3 in 0.1 increments. An appropriate clustering resolution was subsequently chosen guided by silhouette score and biological meaning of clusters. Cluster analysis was done using PrepSCTFindMarkers using the SCT assay and default parameters followed by FindAllMarkers with the only.pos = TRUE, min.pct = 0.10, logfc.threshold = 0.25 and assay = “SCT”. Clusters were annotated based on known marker gene expression. The clustering and cluster analysis of the ventricular zone and white matter subset was performed using the same functions and parameter settings. The white matter subset was exported as a .5had object using the functions SaveH5Seurat and Convert.

#### scVelo

Trajectory analysis was performed using scVelo (Bergen et al., 2020; La Manno et al., 2018) (v0.3.1). Filtering and normalisation were done using pp.filter_and_normalize with default parameters expect min_shared_counts=20, n_top_genes=5000 and moments calculated using pp.moments with default parameters expect n_pcs=30, n_neighbors=30. Thereafter, splicing kinetics was computed using tl.recover_dynamics with default parameters, and gene velocities were estimated using tl.velocity with mode=’deterministic’, use_latent_time=True. Lastly, the velocity graph was computed using tl.velocity_graph with default parameters expect mode_neighbors=’connectivities’ and terminal states and latent time with tl.terminal_states and scv.tl.latent_time, respectively, with default parameters. Trajectories were visualised using pl.velocity_embedding_stream using clusters generated using Seurat.

#### pySCENIC

*In silico* gene regulatory network reconstruction was performed using pySCENIC (Van de Sande et al., 2020) (v0.12.1). A loom file was produced from the .5had output of the Seurat analysis. The GRNBoost2 network inference algorithm grn was run using default parameters and using the mouse or human database of transcription factors for mouse and human data, respectively. Enriched motifs were computed using ctx with default parameters using the mm10 motif ranking database and the v10 annotation database. Regulons were then calculated based on the enriched motifs using df2regulon with default parameters. Lastly, regulon enrichment was calculated using aucell using default parameters and regulon specificity scores were calculated by regulon_specificity_scores with default parameters. Due to the stochastic nature of the network inference algorithm, the analysis was performed 10 times to ensure reproducible results. All databases used are available here: https://resources.aertslab.org/cistarget/.

#### CellOracle

The subset containing gliogenic and neurogenic NEPs from the WM was exported from R into Python and Scanpy (Wolf et al., 2018) (v1.9.8) was used for pseudotime calculation and the generation of an anndata object. *In silico* TF perturbations were carried out using CellOracle (Kamimoto et al., 2023) (v0.16.0) following the standard pipeline using default settings at all functions (https://morris-lab.github.io/CellOracle.documentation/tutorials/simulation.html) and using CellOracle’s pre-computed base GRN from mouse scATAC-seq atlas. The cell TAGACTGTCCAGTTCC from W4 was chosen as the root cell to compute pseudotime. The impact of *in silico* knock-out and over-expression of candidate TFs identified using pySCENIC or from prior literature on cell state transitions was simulated. To simulate the knock-out of a given TF, its expression value was set to 0. For over-expression simulations, the expression was set to the maximum expression value of that TF in any cell.

#### Cell2Fate

To explore the underlying biology resulting in the predicted cell trajectories Cell2Fate (v0.1a0) (Aivazidis et al., 2025) was used. The .5had object from the Seurat analysis is imported as an anndata object and registered using scvi-tools function Cell2fate_DynamicalModel.setup_anndata. The maximal number of modules was calculated using c2f.utils.get_max_modules with default parameters and the model was initialised using c2f.Cell2fate_DynamicalModel with the computed maximum modules and default parameters. Then the model was trained using the function train with default parameters. Following the training, the model was exported using export_posterior with num_samples”: 20, “batch_size” : 2000, “use_gpu” : False, ‘return_samples’: True. The module activity was then computed using compute_module_summary_statistics with default parameters and visualised using plot_module_summary_statistics. Genes enriched in each module were computed and ranked using get_module_top_features using p_adj_cutoff=0.01 and all expressed genes as background. The ranked list of the top 200 most enriched genes in each module was subsequently used for GO term analysis. GO term analysis was performed using PANTHER (Thomas et al., 2022) (v19.0) via the online GUI available at https://pantherdb.org/. GO term analysis for biological process, molecular function and cellular component terms was performed individually with default parameters using Fischer’s Exact test and a false discovery rate calculation to correct for multiple testing.

### Statistical Analysis

Prism (GraphPad) was used for statistical analysis. All data is presented as mean ± s.e.m. Sample sizes for biological replicates are mentioned in text and figure legends where relevant. Statistical tests performed for each analysis were mentioned in the figure legends and Table S7. When one or two-way ANOVA was used, multiple comparisons tests were performed using Tukey’s multiple comparisons test except for Fig. 6K (Šídák’s) and S2A-H (Benjamini, Kreiger and Yukutieli) based on the Prism’s recommendations. See Table S7 for a summary of all statistics performed.

## Data and code availability

The scRNA-seq data is available at ArrayExpress (Accession E-MTAB-13353). The targeted DamID sequencing data is available at NCBI’s Gene Expression Omnibus (GEO Series accession GSE292043). Scripts to process DamID data are available on the following GitHub: https://github.com/AHBrand-Lab/DamID_scripts. Scripts used for data analysis are available on the following GitHub: https://github.com/BayinLab/Christensen_et_al_2025.

## Material availability

Plasmids used are available from the lead contact, N. Sumru Bayin, with a completed Materials Transfer Agreement.

## Acknowledgements

We would like to thank the members of the Bayin Lab for their constructive feedback and Dr James Bae and Niamh Divers for editing the manuscript. We would like to thank Steven Lisgo, Nita Solanky and HDBR, and Roger Baker, Xiaoling He and Cambridge Brain Repair Centre for sharing human fetal tissue. We would like to thank Drs Iva Tchasovnikarova, Emma Rawlins and Julie Ahringer for sharing resources and feedback. Finally, we would like to thank the UBS Gurdon Institute Animal Facility, Gurdon Institute Imaging, Sequencing and Bioinformatics Facilities for their outstanding technical help.

## Funding

We would like to thank our funding sources for supporting our research. JBC: University of Cambridge School of Biological Sciences DTP PhD Studentship and Peter and Emma Thomsen’s Scholarship [1051], GV: Brain Research UK PhD Studentship [PhD23-100024], OAB: Wellcome Trust grants [206194] and [220540/Z/20/A], NSB: Wellcome Career Development Award [227294/Z/23/Z], Royal Society [RGS\R1\231143], Cambridge Stem Cell Institute Seed Funding), AHB: Wellcome Trust Senior Investigator Award [103792], Wellcome Investigator Award [223111], Royal Society Darwin Trust Research Professorship [RP150061]. Gurdon Institute was funded by Wellcome Core Grant [203144] and CRUK Grant [C6946/A24843].

## Author contribution

Conceptualisation: JBC and NSB; Formal analysis: JBC and NSB; Investigation: JBC, APAD, MM, GV, MH, JL; Resources: OB, AHB; Data curation: AJR; Writing – Original draft: JBC and NSB; Writing – Review and edit: JBC, APAD, MM, GV, MH, AJR, JL, OB, AHB, NSB; Visualisation: JBC and NSB; Supervision: AJR, NSB; Project administration: JBC and NSB; Funding acquisition: AHB and NSB.

## Declaration of interests

The authors declare no competing interests.

## Notes

### Competing Interest Statement

The authors have declared no competing interest.

