## Supplementary Figures and Tables for "A conserved differentiation program facilitates inhibitory neuron production in the developing mouse and human cerebellum"

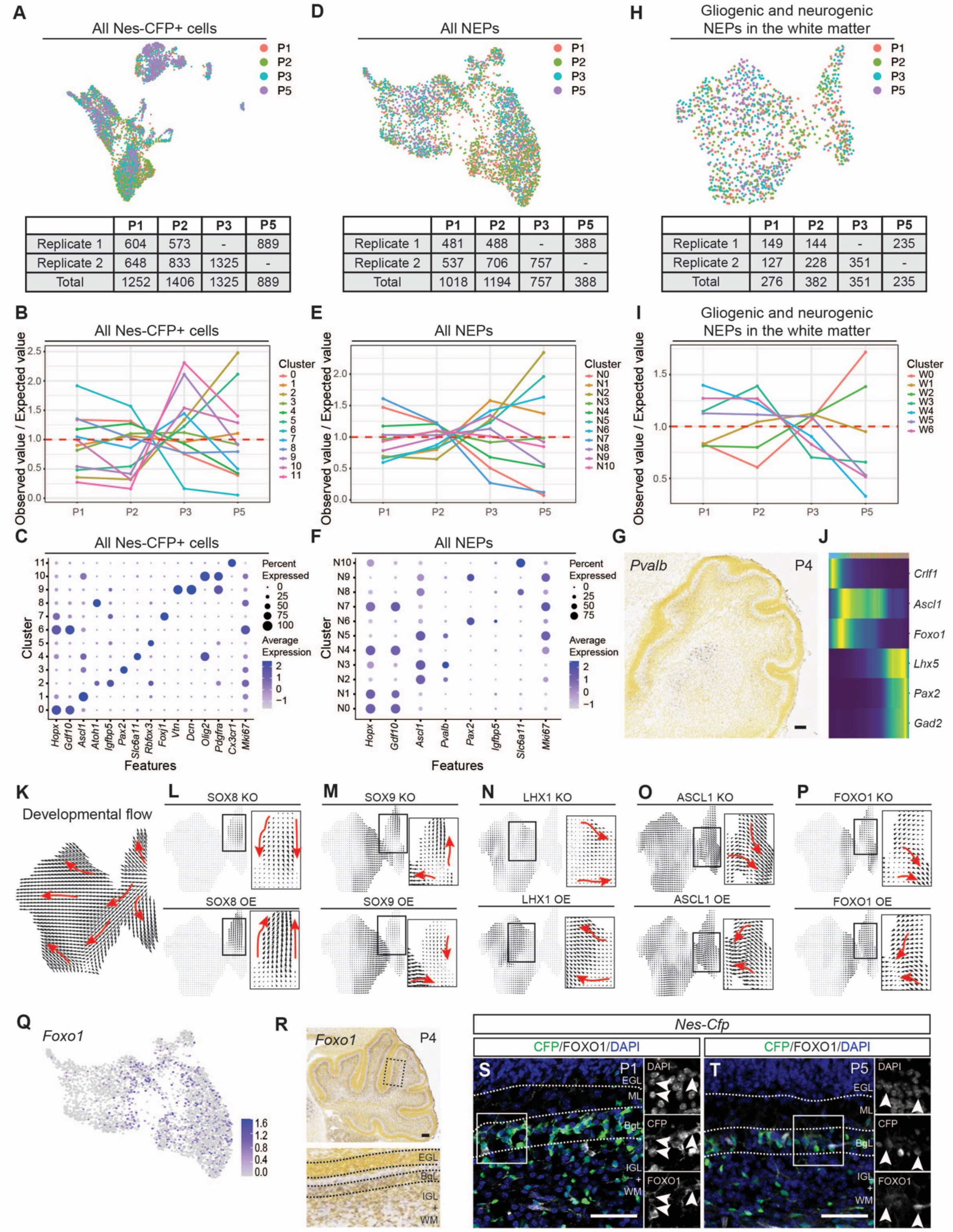


| | P1 | P2 | P3 | P5 |
|---|---|---|---|---|
| Replicate 1 | 604 | 573 | - | 889 |
| Replicate 2 | 648 | 833 | 1325 | - |
| Total | 1252 | 1406 | 1325 | 889 |

| | P1 | P2 | P3 | P5 |
|---|---|---|---|---|
| Replicate 1 | 481 | 488 | - | 388 |
| Replicate 2 | 537 | 706 | 757 | - |
| Total | 1018 | 1194 | 757 | 388 |

| | P1 | P2 | P3 | P5 |
|---|---|---|---|---|
| Replicate 1 | 149 | 144 | - | 235 |
| Replicate 2 | 127 | 228 | 351 | - |
| Total | 276 | 382 | 351 | 235 |

**Fig. S1. Analyses of scRNA-seq of NEPs identify *Foxo1* as a possible regulator of NEP-to-inhibitory neuron differentiation *in vivo*.**

(A-C) Uniform Manifold Approximation and Projection (UMAP) visualisation of all Nes-CFP$^+$ cells coloured by sample timepoint and a table with the number of cells from each replicate at each timepoint (A). Distribution of cells from different timepoints within clusters from Nes-CFP$^+$ cells. The y-axis shows the observed number of cells divided by the expected number of cells. The dashed line indicates a ratio of 1 (B). Dotplot of established lineage markers used to annotate clusters in the Nes-CFP$^+$ subset. The colour gradient signifies the normalised expression. The size of the circle indicates the percentage of cells expressing the specific marker within a cluster (C). (D-F) Similar to (A-C) but performed on the subset containing all NEPs. (G) Allen Brain Atlas P4 RNA *in situ* hybridisation of *Pvalb*. (H-I) Similar to (A-B) but performed on the subset containing only the gliogenic- and neurogenic-NEPs in the white matter. (J) Pseudotemporal ordering of cells from cluster W0-5 coloured by the row-wise normalised expression of different genes. (K) Cellular trajectories are computed by CellOracle using the non-perturbed data. The root cell was chosen from W4. (L-P) CellOracle simulation of either *in silico* knockout or overexpression of SOX8 (L), SOX9 (M), LHX1 (N), ASCL1 (O) and FOXO1 (P). The effect on subpopulations is enlarged in insets. Red arrows are added for visual guidance. (Q) UMAP of all NEPs coloured by normalised expression of *Foxo1*. (R) Allen Brain Atlas P4 RNA *in situ* hybridisation of *Foxo1*. High magnification insets of the expression in the lobules are shown in the bottom panel. (S-T) Immunofluorescent analysis of CFP and FOXO1 on P1 (S) and P5 (T) *Nes-Cfp*/+ mice. Arrowheads indicate CFP$^+$ cells with cytoplasmic FOXO1 expression. EGL: external granule layer, IGL: internal granule layer, ML: molecular layer, BgL: Bergmann glia layer, WM: white matter. Scale bars: 100 µM, except for S-T (50 µM).

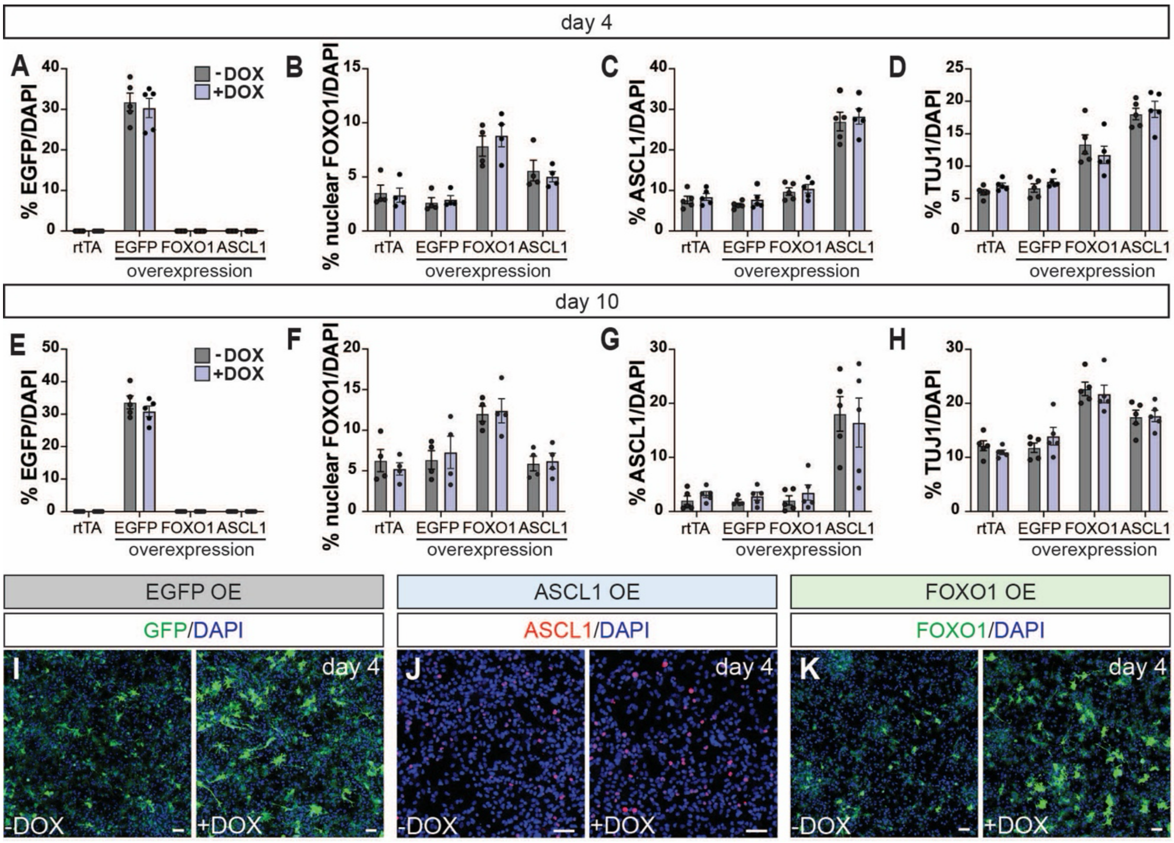


**Fig. S2. The tetracycline-inducible promoter is leaky in primary NEPs.**
(A-H) Quantification of EGFP$^+$, nuclear FOXO1$^+$, ASCL1$^+$ and TUJ1$^+$ cells at day 4 (A-D) and day 10 (E-H) of differentiation in rtTA-NEPs or rtTA-NEPs overexpressing either EGFP, FOXO1 or ASCL1, with and without doxycycline (DOX) treatment. (Multiple Wilcoxon tests, n=5 (except for B and F, n = 4), adjusting for multiple comparisons using the two-stage step-up method of Benjamini, Kreiger and Yukutieli). (I-K) Immunofluorescent analysis of NEPs overexpressing EGFP (EGFP OE) (I), ASCL1 (ASCL1 OE) (J) or FOXO1 (FOXO1 OE) (K) on day 4 of differentiation. Representative images are shown. Graphs show mean ± s.e.m. Scale bars: 50 μm.

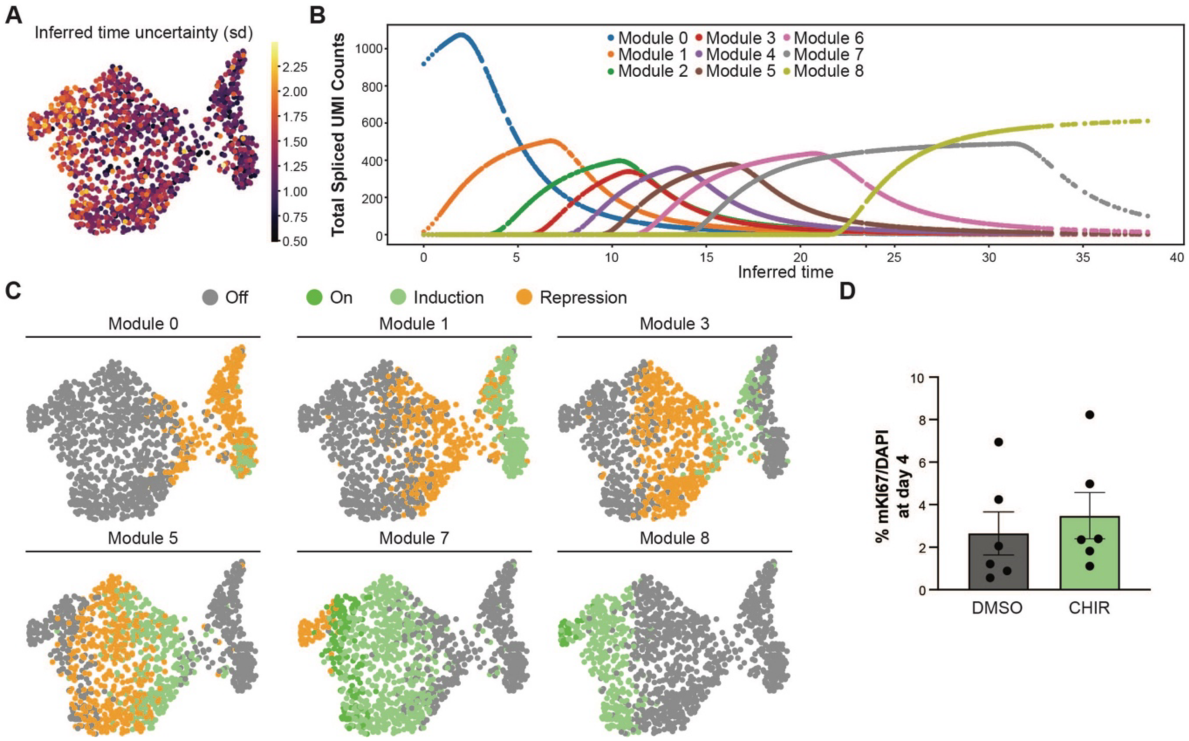


**Fig. S3. Cell2Fate captures sequentially activated modules during NEP-to-inhibitory neuron differentiation.**

(A) UMAP of WM NEPs labelled by inferred time uncertainty computed using Cell2Fate. (B) Total spliced UMI counts from each Cell2Fate module with respect to the computed inferred time. (C) UMAPs of WM NEPs labelled by the module state (Off, On, Induction, Repression) of modules 0, 1, 3, 5, 7 and 8. (D) Quantification of mKI67$^{+}$ cells on cultures treated with either DMSO or CHIR at day 4 of differentiation (ratio paired Student's t-test, n=6). Graphs show mean ± s.e.m.

List of Tables

**Table S1.** The number of cells in each condition, the marker genes used for cluster annotations and the lists of differentially expressed genes in each cluster were identified during the iterative subclustering of scRNA-seq data.

**Table S2.** Regulon specificity score for individual pySCENIC iterations (n=10) and summary of top 3 regulons and the frequency that they were identified.

**Table S3.** Lists of motifs enriched in targeted DamID data that were identified by HOMER.

**Table S4.** List of genes with significant peaks identified by targeted DamID and GO term analysis of those genes that were DamID data.

**Table S5.** Top 200 genes in each Cell2Fate module and GO term analysis using those genes.

**Table S6.** List of primary and secondary antibodies used for immunofluorescent analyses.

**Table S7.** Summary of the statistical comparisons.

**Table S6.** List of primary and secondary antibodies used for immunofluorescent analyses.

| Primary antibodies | | |
|---|---|---|
| Antibody | Catalogue number, supplier | Dilution |
| Rabbit anti-FOXO1 | #2880, Cell Signaling | 1:200 |
| Mouse anti-ASCL1 | 556604, BD Bioscience | 1:500/1:333 |
| Rabbit anti-TUJ1 | ab18207-100ug, Abcam | 1:1000 |
| Chicken anti-GFP | ab13970, Abcam | 1:2000 |
| Goat anti-SOX9 | #AF3075, R&D Systems | 1:500 |
| Goat anti-GAD1 | #AF2086, R&D Systems | 1:250 |
| Mouse anti-GAD2 | AB_2314499, DSHB | 1:100 |
| Mouse anti-PVALB | 195004, Synaptic Systems | 1:500 |
| Rabbit anti-HOPX | HPA030180-100ul, Merck | 1:1000 |
| Mouse anti-MKI67 | ab279653, Abcam | 1:500 |
| Goat anti-SOX2 | AF2018, R&D Systems | 1:1000 |
| Chicken anti-GFAP | Ab4674-50ul, Abcam | 1:1000 |
| **Secondary antibodies** | | |
| Antibody | Catalogue number, supplier | Dilution |
| Donkey anti-rabbit Alexa Fluor™ Plus 488 | #32790, ThermoFischer Scientific | 1:500 |
| Donkey anti-mouse Alexa Fluor™ Plus 555 | #32773, ThermoFischer Scientific | 1:500 |
| Donkey anti-goat Alexa Fluor™ Plus 488 | 15930877, FisherScientific | 1:500 |
| Donkey anti-mouse Alexa Fluor™ Plus 647 | 15927745, FisherScientific | 1:500 |
| Donkey anti-chicken Alexa Fluor™ Plus 488 | 17777517, Fisher | 1:500 |
| Donkey anti-rabbit Alexa Fluor™ Plus 647 | 16239260, FisherScientific | 1:500 |
| Donkey anti-rabbit Alexa Fluor™ Plus 555 | A32794, ThermoFisher | 1:500 |
| Donkey anti-mouse Alexa Fluor™ Plus 488 | #32790, ThermoFischer Scientific | 1:500 |
| Streptavadin, Alexa Fluor™ 555 Conjugate | S32355, ThermoFisher | 1:500 |
| DAPI | MBD0015-1ML, Sigma-Aldrich | 1:1000 |

**Table S7**. Summary of the statistical comparisons.

| Figure | Test Performed | p-value | Multiple comparisons | |
|---|---|---|---|---|
| Figure 2F | Ordinary one-way ANOVA | $F_{(5, 46)}$ = 9. 439 P<0.0001 | Tuckey's multiple comparisons | |
| | | | Comparisons | Adjusted p value |
| | | | 0 vs 2 | 0.9909 |
| | | | 0 vs 4 | 0.8883 |
| | | | 0 vs 6 | 0.0726 |
| | | | 0 vs 10 | <0.0001 |
| | | | 0 vs 14 | 0.0004 |
| | | | 2 vs 4 | 0.9966 |
| | | | 2 vs 6 | 0.2474 |
| | | | 2 vs 10 | 0.0004 |
| | | | 2 vs 14 | 0.0027 |
| | | | 4 vs 6 | 0.5131 |
| | | | 4 vs 10 | 0.0017 |
| | | | 4 vs 14 | 0.011 |
| | | | 6 vs 10 | 0.1287 |
| | | | 6 vs 14 | 0.4701 |
| | | | 10 vs 14 | 0.9526 |
| Figure 2K | Ordinary one-way ANOVA | $F_{(5, 46)}$ = 7. 720 P<0.0001 | Tuckey's multiple comparisons | |
| | | | Comparisons | Adjusted p value |
| | | | 0 vs 2 | 0.0689 |
| | | | 0 vs 4 | 0.9948 |
| | | | 0 vs 6 | 0.9986 |
| | | | 0 vs 10 | 0.2087 |
| | | | 0 vs 14 | 0.0938 |
| | | | 2 vs 4 | 0.0182 |
| | | | 2 vs 6 | 0.0257 |
| | | | 2 vs 10 | 0.0001 |
| | | | 2 vs 14 | <0.0001 |
| | | | 4 vs 6 | >0.9999 |
| | | | 4 vs 10 | 0.4615 |
| | | | 4 vs 14 | 0.2678 |
| | | | 6 vs 10 | 0.3891 |
| | | | 6 vs 14 | 0.212 |
| | | | 10 vs 14 | >0.9999 |
| Figure 2L | Ordinary one-way ANOVA | $F_{(5, 46)}$ = 3. 575 P=0.0082 | Tuckey's multiple comparisons | |
| | | | Comparisons | Adjusted p value |
| | | | 0 vs 2 | >0.9999 |
| | | | 0 vs 4 | 0.1229 |
| | | | 0 vs 6 | 0.0286 |
| | | | 0 vs 10 | 0.2105 |
| | | | 0 vs 14 | 0.9106 |

<table>
<tr><td></td><td></td><td></td><td><table>
<tr><td>2 vs 4</td><td>0.1843</td></tr>
<tr><td>2 vs 6</td><td>0.0474</td></tr>
<tr><td>2 vs 10</td><td>0.293</td></tr>
<tr><td>2 vs 14</td><td>0.9632</td></tr>
<tr><td>4 vs 6</td><td>0.9895</td></tr>
<tr><td>4 vs 10</td><td>>0.9999</td></tr>
<tr><td>4 vs 14</td><td>0.6247</td></tr>
<tr><td>6 vs 10</td><td>0.9837</td></tr>
<tr><td>6 vs 14</td><td>0.267</td></tr>
<tr><td>10 vs 14</td><td>0.749</td></tr>
</table></td></tr>
<tr><td>Figure 3C</td><td>Row matched one-way ANOVA</td><td>$F_{(1,4)}$ = 149.0<br>p-value = 0.0003</td><td>Tukey’s multiple comparisons comparing the mean of every column.<br>EGFP OE vs FOXO1 OE : adjusted p-value = 0.0006<br>EGFP OE vs ASCL1 OE : adjusted p-value = 0.0006<br>FOXO1 OE vs ASCL1 OE : adjusted p-value = NA</td></tr>
<tr><td>Figure 3E</td><td>Row matched one-way ANOVA</td><td>$F_{(1.332,5.328)}$ = 138.6<br>p-value = <0.0001</td><td>Tukey’s multiple comparisons comparing the mean of every column.<br>EGFP OE vs FOXO1 OE : adjusted p-value = 0.0501<br>EGFP OE vs ASCL1 OE : adjusted p-value = 0.0005<br>FOXO1 OE vs ASCL1 OE : adjusted p-value = 0.0006</td></tr>
<tr><td>Figure 3G</td><td>Row matched one-way ANOVA</td><td>$F_{(1.024,3.073)}$ = 35.83<br>p-value = 0.0087</td><td>Tukey’s multiple comparisons comparing the mean of every column.<br>EGFP OE vs FOXO1 OE : adjusted p-value = 0.0172<br>EGFP OE vs ASCL1 OE : adjusted p-value = 0.0072<br>FOXO1 OE vs ASCL1 OE : adjusted p-value = 0.0278</td></tr>
<tr><td>Figure 3J</td><td>Row matched one-way ANOVA</td><td>$F_{(1.722,6.889)}$ = 39.25<br>p-value = 0.0002</td><td>Tukey’s multiple comparisons comparing the mean of every column.<br>EGFP OE vs FOXO1 OE : adjusted p-value =</td></tr>
</table>

| | | | |
|---|---|---|---|
| | | | 0.00319<br>EGFP OE vs ASCL1 OE : adjusted p-value = 0.0024<br>FOXO1 OE vs ASCL1 OE : adjusted p-value = 0.0184 |
| Figure 3K | Row matched one-way ANOVA | $F_{(1.472,5.888)}$ = 17.37<br>p-value = 0.0044 | Tukey's multiple comparisons comparing the mean of every column.<br>EGFP OE vs FOXO1 OE : adjusted p-value = 0.0017<br>EGFP OE vs ASCL1 OE : adjusted p-value = 0.1287<br>FOXO1 OE vs ASCL1 OE : adjusted p-value = 0.1207 |
| Figure 3N | Ratio paired t-test | Scrambled vs shRNA1 : p-value = 0.0217<br>Scrambled vs shRNA2 : p-value = 0.0124 | |
| Figure 3P | Ratio paired t-test | Scrambled vs shRNA1 : p-value = 0.0607<br>Scrambled vs shRNA2 : p-value = 0.044 | |
| Figure 3Q | Ratio paired t-test | Scrambled vs shRNA1 : p-value = 0.9153<br>Scrambled vs shRNA2 : p-value = 0.8407 | |
| Figure S2A | Multiple Wilcoxon tests between control and doxycycline conditions, adjusting p-value using the two- | | EGFP : q-value = 0.820625 |

| | | | |
|---|---|---|---|
| | stage step-up method of Benjamini, Kreiger and Yukutieli. | | |
| Figure S2B | Multiple Wilcoxon tests between control and doxycycline conditions, adjusting p-value using the two-stage step-up method of Benjamini, Kreiger and Yukutieli. | | rtTA : q-value = 0.75750<br>EGFP OE: q-value = 0.50500<br>FOXO1 OE: q-value = 0.883750<br>ASCL1 OE : q-value = 0.841667 |
| Figure S2C | Multiple Wilcoxon tests between control and doxycycline conditions, adjusting p-value using the two-stage step-up method of Benjamini, Kreiger and Yukutieli. | | rtTA : q-value = 0.820625<br>EGFP OE: q-value = 0. 820625<br>FOXO1 OE: q-value = 0. 820625<br>ASCL1 OE : q-value = 0. 820625 |
| Figure S2D | Multiple Wilcoxon tests between control and doxycycline conditions, adjusting p-value using the two-stage step-up method of Benjamini, Kreiger and Yukutieli. | | rtTA : q-value = 0.252500<br>EGFP OE: q-value = 0. 252500<br>FOXO1 OE: q-value = 0. 252500<br>ASCL1 OE : q-value = 0. 631250 |
| Figure S2E | Multiple Wilcoxon tests between control and doxycycline conditions, adjusting p-value using the two-stage step-up method of Benjamini, Kreiger and Yukutieli. | | EGFP control vs EGFP DOX: q-value = 0.063125 |
| Figure S2F | Multiple Wilcoxon tests between | | rtTA : q-value = 0.883750 |

| | | | |
|---|---|---|---|
| | control and doxycycline conditions, adjusting p-value using the two-stage step-up method of Benjamini, Kreiger and Yukutieli. | | EGFP OE: q-value = 0. 883750<br>FOXO1 OE: q-value = 0.883750<br>ASCL1 OE : q-value = 0. 883750 |
| Figure S2G | Multiple Wilcoxon tests between control and doxycycline conditions, adjusting p-value using the two-stage step-up method of Benjamini, Kreiger and Yukutieli. | | rtTA : q-value = 0.25250<br>EGFP OE: q-value = 0. 441875<br>FOXO1 OE: q-value = 0. 441875<br>ASCL1 OE : q-value = 0. 441875 |
| Figure S2H | Multiple Wilcoxon tests between control and doxycycline conditions, adjusting p-value using the two-stage step-up method of Benjamini, Kreiger and Yukutieli. | | rtTA : q-value = 0.589157<br>EGFP OE: q-value = 0. 589157<br>FOXO1 OE: q-value = 0. 589157<br>ASCL1 OE : q-value > 0. 999999 |
| Figure 4K | Row matched one-way ANOVA | $F_{(1.245, 3.734)}$ = 46.00<br>p-value = 0.0029 | Tukey's multiple comparisons comparing the mean of every column.<br>EGFP OE vs FOXO1 OE : adjusted p-value = 0.0845<br>EGFP OE vs ASCL1 OE : adjusted p-value = 0.0177<br>FOXO1 OE vs ASCL1 OE : adjusted p-value = 0.0090 |
| Figure 4M | Ordinary one-way ANOVA | $F_{(2, 9)}$ = 5.475<br>p-value = 0.0278 | Tukey's multiple comparisons comparing the mean of every column.<br>EGFP OE vs FOXO1 OE : adjusted p-value = 0.0225 |

<table>
<tr><td></td><td></td><td></td><td>EGFP OE vs ASCL1 OE : adjusted p-value = 0.3198<br>FOXO1 OE vs ASCL1 OE : adjusted p-value = 0.2339</td></tr>
<tr><td>Figure 4P</td><td>Paired t-test</td><td>p-value = 0.0063</td><td></td></tr>
<tr><td>Figure 5L</td><td>Ratio paired t-test</td><td>p-value = 0.0016</td><td></td></tr>
<tr><td>Figure 5M</td><td>Ratio paired t-test</td><td>p-value = 0.1677</td><td></td></tr>
<tr><td>Figure 5P</td><td>Ratio paired t-test</td><td>p-value = 0.0013</td><td></td></tr>
<tr><td>Figure S5C</td><td>Ratio paired t-test</td><td>p-value = 0.1208</td><td></td></tr>
<tr><td>Figure 6I</td><td>Row matched one-way ANOVA</td><td>$F_{(2.919, 23.35)}$ = 6.614<br>p-value = 0.023</td><td>Tuckey’s multiple comparisons
<table>
<tr><th>Comparisons</th><th>Adjusted p value</th></tr>
<tr><td>0 vs 2</td><td>0.7433</td></tr>
<tr><td>0 vs 4</td><td>0.1103</td></tr>
<tr><td>0 vs 6</td><td>0.4186</td></tr>
<tr><td>0 vs 10</td><td>0.3949</td></tr>
<tr><td>2 vs 4</td><td>0.0135</td></tr>
<tr><td>2 vs 6</td><td>0.0176</td></tr>
<tr><td>2 vs 10</td><td>0.0836</td></tr>
<tr><td>4 vs 6</td><td>0.9367</td></tr>
<tr><td>4 vs 10</td><td>0.7635</td></tr>
<tr><td>6 vs 10</td><td>0.9997</td></tr>
</table></td></tr>
<tr><td>Figure 6J</td><td>Row matched one-way ANOVA</td><td>$F_{(2.552, 20.41)}$ = 6.812<br>p-value = 0.0033</td><td>Tuckey’s multiple comparisons
<table>
<tr><th>Comparisons</th><th>Adjusted p value</th></tr>
<tr><td>0 vs 2</td><td>0.0077</td></tr>
<tr><td>0 vs 4</td><td>0.9996</td></tr>
<tr><td>0 vs 6</td><td>>0.9999</td></tr>
<tr><td>0 vs 10</td><td>0.2949</td></tr>
<tr><td>2 vs 4</td><td>0.1994</td></tr>
<tr><td>2 vs 6</td><td>0.1015</td></tr>
<tr><td>2 vs 10</td><td>0.0092</td></tr>
<tr><td>4 vs 6</td><td>0.9976</td></tr>
<tr><td>4 vs 10</td><td>0.5795</td></tr>
<tr><td>6 vs 10</td><td>0.2865</td></tr>
</table></td></tr>
<tr><td>Figure 6K</td><td>Ordinary two-way ANOVA</td><td>Species: $F_{(1, 70)}$ = 11.51, P=0.0011</td><td>Šídák’s multiple comparisions
<table>
<tr><th>Comparisons</th><th>Adjusted p value</th></tr>
<tr><td>0</td><td>0.8899</td></tr>
</table></td></tr>
</table>

<table>
<tr><td></td><td></td><td>Timepoint: $F_{(4, 70)}$ = 1.388, P=0.2471<br>Interaction: $F_{(4, 70)}$ = 1.604, P=0.1829</td><td><table><tr><td>2</td><td>0.0023</td></tr><tr><td>4</td><td>0.9971</td></tr><tr><td>6</td><td>0.5715</td></tr><tr><td>10</td><td>0.7729</td></tr></table></td></tr>
<tr><td>Figure 6O</td><td>Ratio paired t-test</td><td>p-value = 0.0233</td><td></td></tr>
<tr><td>Figure 6P</td><td>Ratio paired t-test</td><td>p-value = 0.2703</td><td></td></tr>
<tr><td>Figure 6S</td><td>Ratio paired t-test</td><td>p-value = 0.0097</td><td></td></tr>
</table>
